# Depletion of Sun1/2 induces heterochromatin accrual in mesenchymal stem cells during adipogenesis

**DOI:** 10.1101/2022.02.15.480528

**Authors:** Matthew Goelzer, Sean Howard, Anamaria Zavala, Daniel Conway, Janet Rubin, Andre J van Wijnen, Gunes Uzer

## Abstract

Critical to the mechano-regulation of mesenchymal stem cells (MSC), Linker of the Nucleoskeleton and Cytoskeleton (LINC) complex transduces cytoskeletal forces to the nuclei. The LINC complex contains outer nuclear membrane Nesprin proteins that associate with the cytoskeleton and their inner nuclear membrane couplers, Sun proteins. In addition to coupling Nesprin-associated cytoskeletal elements to inner nuclear membrane, Sun proteins also function in regulating nuclear mechanics and chromatin tethering to inner nuclear membrane. This suggests that release of LINC-mediated cytoskeletal connections from cell nuclei may have different effects on chromatin organization and MSC differentiation than those due to ablation of intra-nuclear Sun proteins. To test this hypothesis we compared Sun1/2 depletion with expression of the dominant-negative KASH domain (dnKASH) that inhibits Nesprin-Sun association. In cells cultured under adipogenic conditions, disconnecting LINC from the cytoskeleton through dnKASH expression, increased adipogenic gene expression and fat droplet formation; heterochromatin H3K27 and H3K9 methylation was unaltered. In contrast, Sun1/2 depletion inhibited adipogenic gene expression and fat droplet formation; as well the anti-adipogenic effect of Sun1/2 depletion was accompanied by increased H3K9me3, which was enriched on *Adipoq*, silencing this fat locus. We conclude that releasing the nucleus from cytoskeletal constraints via dnKASH accelerates adipogenesis while depletion of Sun1/2 increases heterochromatin accrual on adipogenic genes in a fashion independent of LINC complex function. Therefore, while these two approaches both disable LINC functions, their divergent effects on the epigenetic landscape indicate they cannot be used interchangeably to study mechanical regulation of cell differentiation.

## Introduction

The Linker of the Cytoskeleton and Nucleoskeleton (LINC) complex is composed of Sun and Nesprin proteins^1^. N-termini of Nesprin proteins associate with cytoskeletal filaments such as actin^2,3^, microtubules^4,5^ and other adaptor proteins^6–8^ to provide dynamic nucleo-cytoplasmic connections at the outer nuclear membrane. Nesprin C-termini, located inside the perinuclear space, bind to C-termini of the Sun proteins via their KASH domain^9^. The Sun protein N-termini pierce the inner nuclear membrane to interface with both A- and B-type Lamins^10^, Emerin^11^, and nuclear pore complexes^12^ as well as providing organizational capacity to chromatin by interacting with telomere ends of DNA during DNA repair^13^ and meiosis^14,15^. In this capacity, LINC complexes serve as mechanosensitive adaptors between cytoplasm and nucleus to regulate both physical and biochemical signal transduction to the nucleus^16^.

During development and in mechanically active tissues such as bone and muscle, progenitor cell differentiation is highly dependent upon a cell’s ability to sense extracellular mechanical cues and transmit this information into nucleus and alter chromatin structure. Therefore, disrupting Nesprin-Sun binding of the LINC complex via expression of a dominant-negative KASH (dnKASH) and depletion of either Sun or Nesprin proteins have been used interchangeably to study the effects of nucleo-cytoskeletal connectivity on cell function and differentiation^17^. Indeed, both dnKASH overexpression and Nesprin/Sun depletion approaches have been shown to affect progenitor cell differentiation. dnKASH expression, for example, increases histone deacetylase (HDAC) activity in human MSCs; resulting in decreased expression of the osteogenesis marker Runx2 and increased expression of the adipogenesis marker Pparg^18^. Depletion of both Nesprin2-Giant^20^ and Sun1/2 are associated with epidermal thickening^19,20^, in case of Sun1/2 co-depletion through decreased differentiation of keratinocyte progenitors.

Disabling LINC complex function may affect cell fate via number of mechanisms. Our group and others have shown that both dnKASH expression^21,22^ and Sun1/2 co-depletion^23,24^ alters the force generation and dynamics of focal adhesions. In mesenchymal stem cells (MSC), both approaches impair the activation of signaling molecules FAK (Focal Adhesion Kinase) and Akt (Serine/Threonine Kinase) in response to mechanical vibrations, thus limiting the RhoA mediated increase in cell contractility^25^. LINC-mediated transmission of contractile forces to the nuclear envelope also affect the nuclear entry of mechanotransducer proteins β-Catenin and Yap with proliferative^26–28^ and anti-adipogenic^29–32^ functions in MSC. For example, both dnKASH^33^ expression and Sun1/2 co-depletion^34^ decrease nuclear β-Catenin levels. In the case of Yap, dnKASH expression^35^ and depletion of Nesprin1^36^ both inhibit nuclear translocation through nuclear pore complexes in response to mechanical stimulus, potentially by limiting cytoskeleton-induced nuclear deformations, or by impairing association with Yap’s nuclear transporter, Importin 7^37^. LINC-mediated cytoskeletal connections also affect deformations of the inner nuclear membrane: for example, it dnKASH expression reduces stretching of LaminA/C measured via FRET-based LaminA/C force sensor^38^, and decreases the chromatin deformations during cardiomyocyte contractions^39^. Similarly, stress-induced chromatin stretching, caused by magnetic bead motions bound to cell membrane, is abolished upon depletion of Sun1/2 proteins^40^. Therefore, both depletion of Sun proteins and disrupting LINC connectivity directly affect cell mechanosignaling and chromatin organization.

Another important regulator of cell fate is the nuclear mechanics determined by A-type Lamins and chromatin^41^. LaminA/C, the most well characterized structural protein at the inner nuclear envelope, has been shown to impact differentiation of MSCs^42,43^. LaminA/C modulates cell differentiation by regulating chromatin organization^44,45^ which we have shown to be independent of mechano-signaling in MSCs^46^. Emerging data suggests that depleting Sun1/2 proteins softens nuclei^47^ and results in similar nuclear stiffness measures t comparable cells depleted of LaminA/C depleted^48^. dnKASH expression on the other hand does not significantly alter nuclear mechanics^49^. In fact, we have utilized atomic force microscopy of live isolated cell nuclei to show that dnKASH separation of nucleus and cytoskeletal structure slightly increases the stiffness of isolated nuclei^48^.

Indeed, Sun1/2 elements provide organizational capacity to chromatin through associations with LEM domain proteins (LAP2, Emerin, MAN1 domain), direct links to chromatin^46,47^ and, ultimately, regulation of the genome^50,51^. In mammalian cells, Sun1/2 proteins tether chromatin to the nuclear envelope through direct connections to Emerin^52^. During meiosis, *Sun1^−/−^* mouse cells display disrupted telomere association to nuclear envelope^53^. Chromatin capture studies further show that depletion of Sun proteins disrupt the alignment of different chromosome ends via altering telomere binding to the nuclear envelope^54^. This suggests that disrupting LINC at the level of the outer (dnKASH) and inner (Sun1/2) membrane may result in different dispositions of cell fate beyond LINC complex-mediated cytoskeletal tethering. To test this hypothesis we compared siRNA-mediated depletion of Sun1/2 to the over expression of the dnKASH domain to block Sun-Nesprin association^25,55^ in a model of MSC adipogenesis.

## Results

### siSun and dnKASH expression alter nuclear morphology

As nuclear shape is affected by both in LINC connectivity and nuclear mechanics^56,57^, we quantified the effects of Sun1/2 depletion and dnKASH overexpression on nuclear morphology under growth medium (GM) condition. We first compared MSCs treated with control (siCntl) and Sun1/2 siRNAs (siSun), stained against Sun1 (green), Sun2 (red), and DNA (Hoechst 33342, blue). Shown in **Fig. 1a-f**, siSun treatment reduced Sun1 and Sun2 intensity levels by 47% and 52%, increased nuclear area by 7% and perimeter by 8% while decreasing the nuclear circularity by 9%.

**Figure 1.**
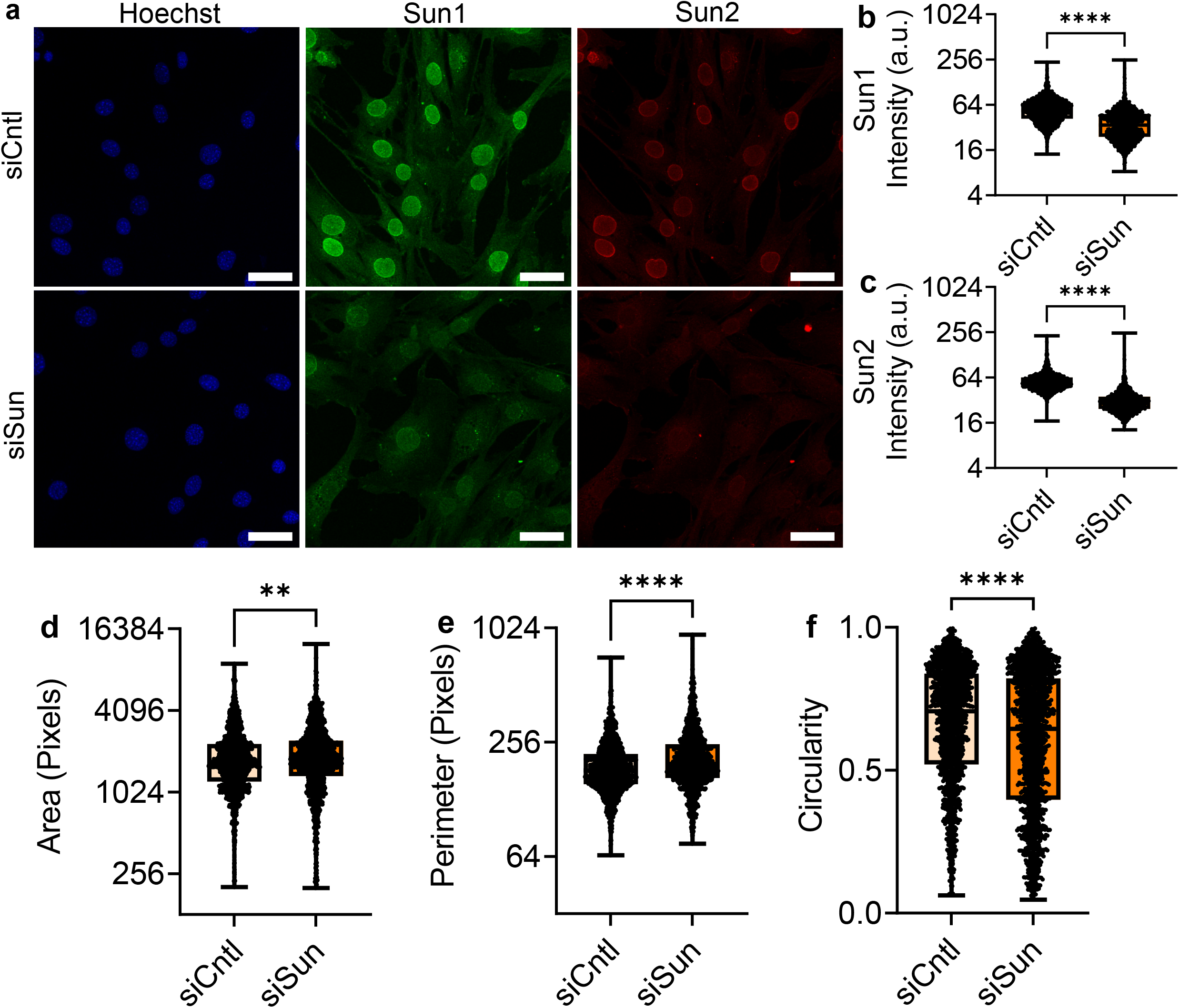
Sun1/2 depletion alters nuclear morphology. **a** Representative images of MSCs treated with siRNA targeting Sun1/2 (siSun) which were stained for Sun1 (green), Sun2 (red), and DNA (blue). **b** siSun treated cells had 47% decrease of Sun1 intensity (n = 1478, P < 0.0001). **c** Sun2 intensity levels were decreased by 52% in siSun treated cells (n = 1478, P < 0.0001). **d** MSCs treated with siSun had an increase in nuclear area by 7% (n = 1478, p 0<0.01). **e** Nucleus perimeter decreased in siSun treated MSCs by 8% (n = 1478, p<0.0001). **f** Nuclear circularity decreased by 9% in siSun treated cells (n = 1478, p<0.001). Comparisons were made against control using non-parametric Mann-Whitney Test where * P < 0.05, ** P < 0.01, *** P < 0.001, **** P < 0.0001. Scale bar represents 50μm.

To disable LINC function, we stably infected MSCs via lentivirus harboring a doxycycline (Dox) inducible mCherry-tagged KASH domain (dnKASH-MSCs). 1 μg/ml Dox was added to cell culture medium to induce mCherry-KASH and remove Nesprins from the nuclear envelope (referred as +Dox, **Fig.S1**). No Dox treatment was used as control. Shown in **Fig.2**, +Dox treatment in GM (GM+Dox) increased mCherry intensity by 133% as well as decreasing nuclear area and circularity by 14% and 6%. Nuclear perimeter was unaltered.

**Figure 2.**
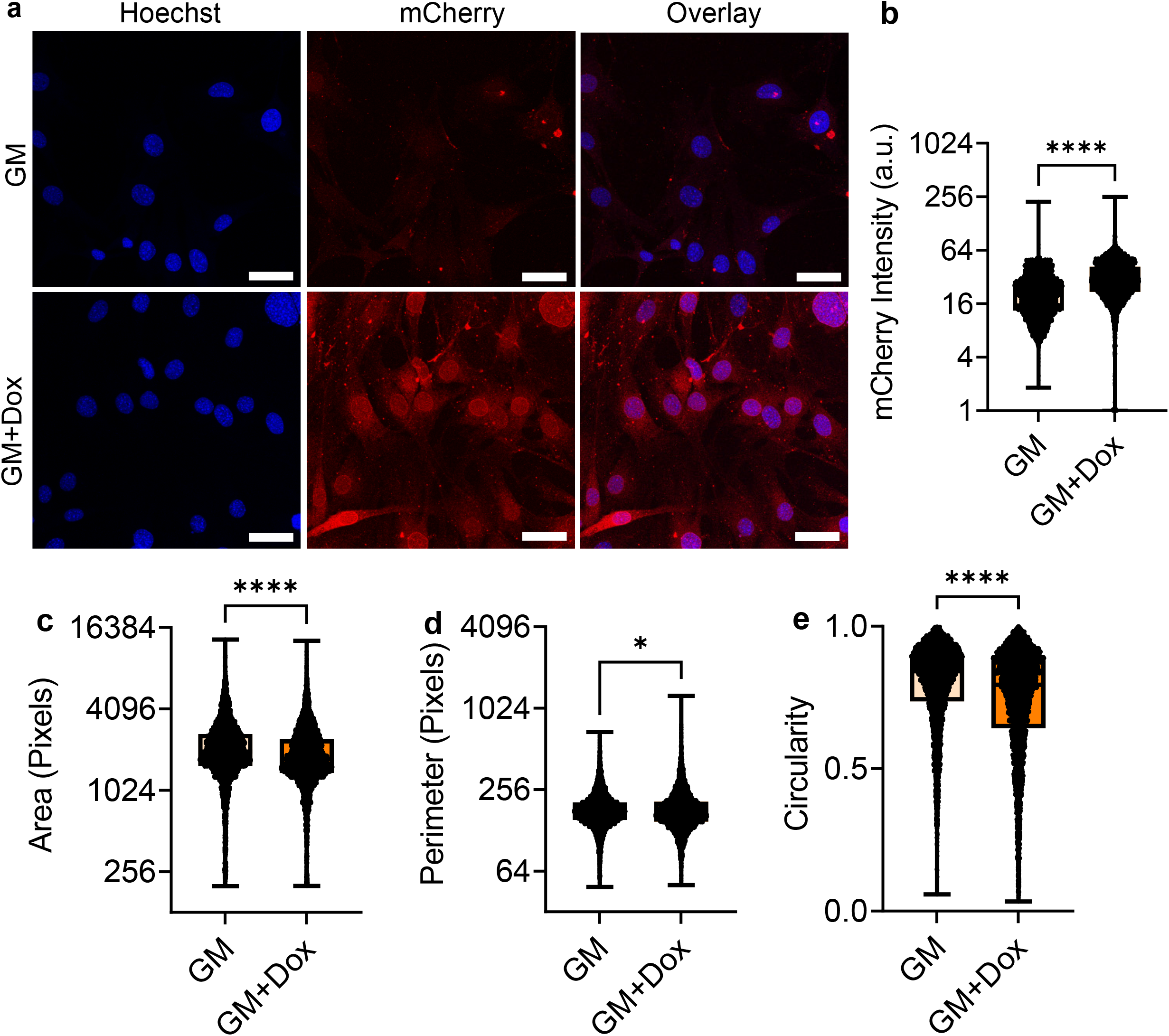
dnKASH expression reduces nuclear area. **a** Representative photos of doxycycline induced DNKASH cells. Images show DNKASH tagged with mCherry (Red) and DNA (Blue). **b** mCherry intensity levels increased by 133% in doxycycline treated MSCs (n = 5332, P < 0.0001). **c** Doxycycline treated MSCs experienced a 14% decrease of nuclear area (n = 5322, p 0.001). **d** Nuclear perimeter had a slight decrease of 1% in doxycycline treatment group (n = 5322, P < 0.05). **e** Nuclear circularity decreased 6% in the doxycycline treatment group (n = 5322, P < 0.0001). Comparisons were made against control using non-parametric Mann-Whitney Test where * P < 0.05, ** P < 0.01, *** P < 0.001, **** P < 0.0001. Scale bar represents 50μm.

### Depletion of Sun1/2 inhibits adipogenesis

To compare adipogenesis of siCntl and siSun treatments, MSCs were cultured under adipogenic differentiation media (AM) for five days and compared to undifferentiated GM controls. Shown in **Fig. 3a-f**, addition of AM onto siCntl MSCs increased adipogenic proteins Adipoq (600%), Cebpa (400%), Pparg (200%) and mean lipid staining per cell by 2000%. In contrast, siSun treated MSCs showed no increase in adipogenic proteins with the exception of Cebpa (185%), and mean lipid staining showed a small 20% increase. This indicated an impaired adipogenic differentiation in siSun treated cells.

**Figure 3:**
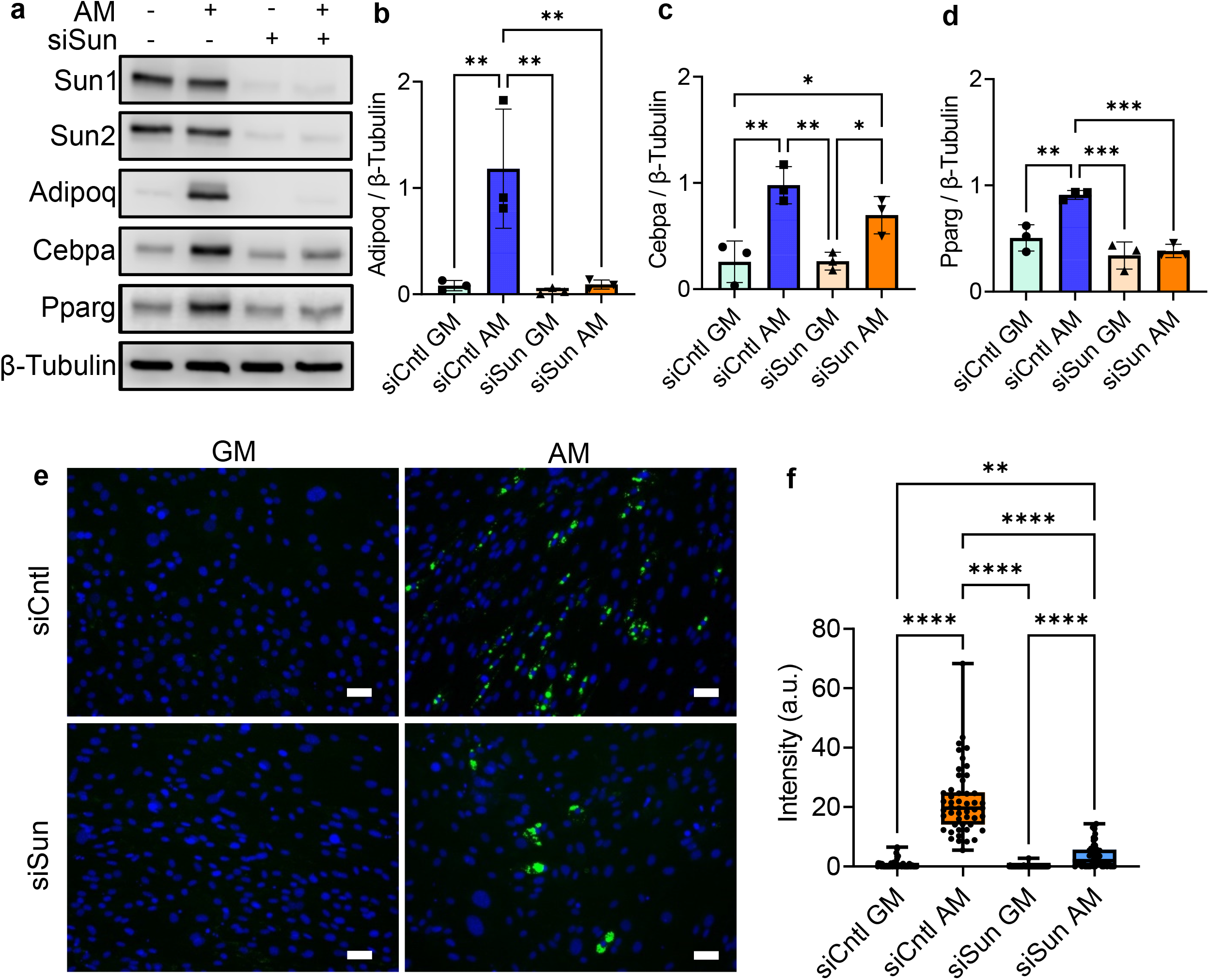
Depletion of Sun1/2 inhibits adipogenesis. **a** Western analysis of adipogenesis markers Adipoq, Cebpa, and Pparg in growth media and adipogenic media during siSun and siCntl treatment. **b** Analysis of Adipoq protein levels. Comparison of adipogenic siSun and siCntl groups showed a 92% reduction of Adipoq (n = 3, P < 0.01). **c** Cebpa experienced a non-significant reduction of 38% in protein levels in siSun cells compared to siCntl cells during adipogensis (n = 3, p = 0.22). **d** Pparg levels decreased by 58% in adipogenic siSun compared to siCntl (n = 3, P < 0.001). **e** Representative images of lipid droplet florescence images where MSCs are stained for lipid droplets (green) and DNA (blue). **f** Quantification of the mean florescent lipid droplet intensity per cell from individual imaging fields shows a significant reduction of 83% in lipid droplet amounts in adipogenic siSun treatment compared to siCntl (n = 50, P < 0.0001). Western analysis group comparisons were made using One-Way ANOVA. Lipid droplet intenisty group comparisons were made using Kruskal-Wallis test. * P < 0.05, ** P < 0.01, *** P < 0.001, **** P < 0.0001. Scale bar represents 50μm.

We next performed RNA-seq analysis on siSun and siCntl samples cultured under GM and AM conditions. We used DESEQ2 analyses to filter genes with significant expression differential between treatment pairs (fold change ≥ 2-fold, P < 0.05). Hierarchical mapping of these differentials (**Fig. 4a**) showed clustering of siSun and siCntl treatments with further sub-clustering of AM and GM conditions within each treatment. Shown in **Fig. 4b**, variance of two principle components were 16.2% and 19.9%. Comparing the gene profiles between AM treated siCntl and siSun groups, DAVID and STRING analyses identified thirteen differentially expressed pathways with an FDR < 0.05 that were either significantly up regulated (**Figs. 4c** **& S2**) and down regulated (**Figs. 4d** **& S3**). Sun1/2 depletion significantly decreased the expression of adipogenic genes Adipoq, Fabp4, Lipe, Plin1, Cidec, Agpath2, Acsl1 and mgll genes (**Fig.4e**) and down-regulated three adipogenesis or lipid metabolism related pathways (**Fig. 4d**, blue bars). Inflammatory response was also robustly upregulated in the siSun group, potentially highlighting a regulatory role of Sun1/2 for inflammation pathways (**Fig.4e**). Tables of DAVID pathway analysis can be found in **Tables S1** and **S2**.

**Figure 4:**
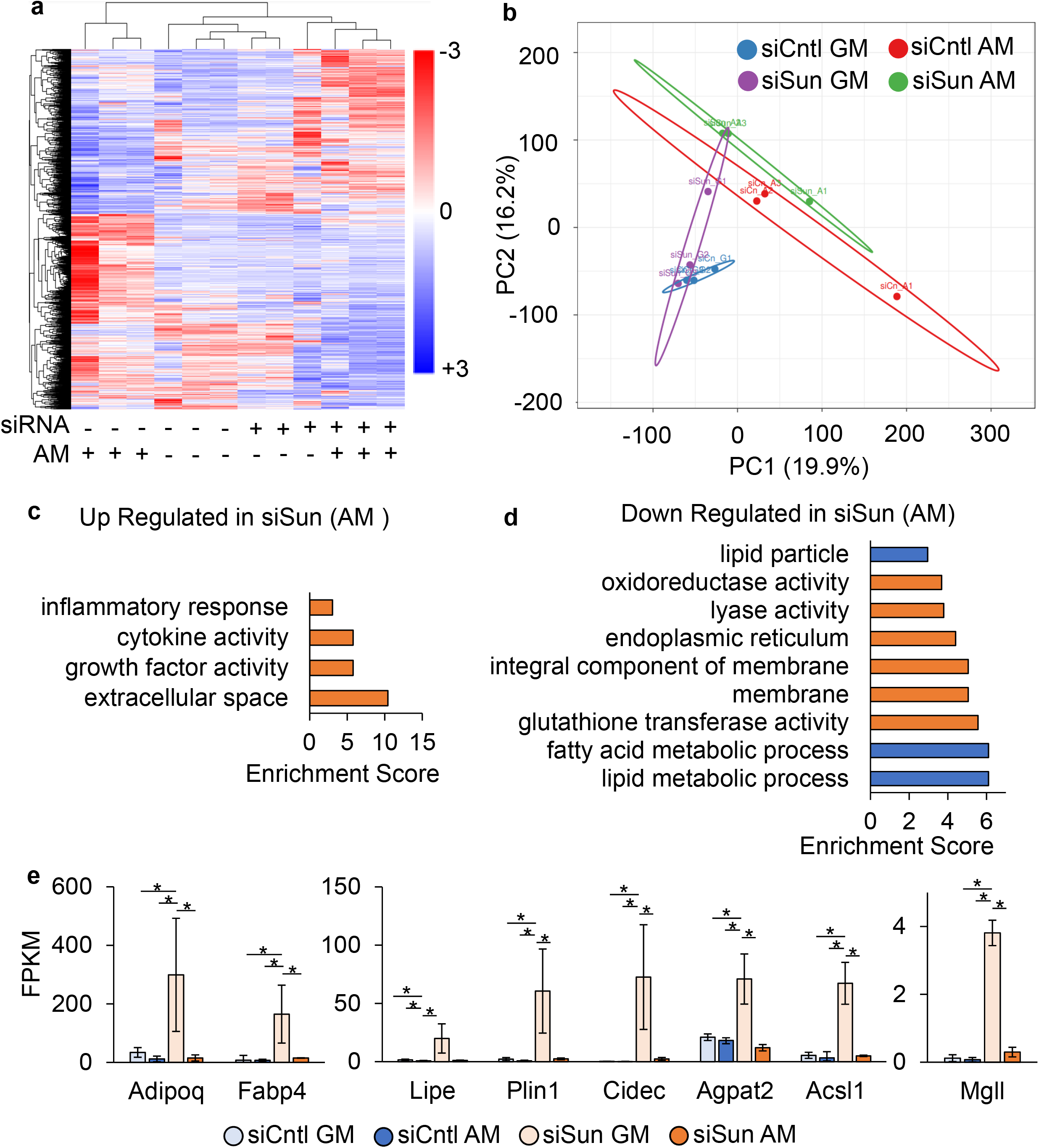
Sun1/2 depletion decreases adipogenesis and lipid metabolism related genes. **a** Heatmap of genes with significant differential (FC > 1 and P < 0.05) gene expression during Sun1/2 depletion (n = 24). **b** Principle component plot where principal component 1 and principal component 2 explain 19.9% and 16.2% of the total variance, respectively. Prediction ellipses indicate that with a probability of 0.95, a new observation from the same group will fall inside the ellipse. *n* = 24 data points. **c** DAVID analysis of genes up regulated in siSun treatment compared to siCntl. Pathways selected have FDR < 0.05. **d** DAVID analysis of genes down regulated in siSun group compared to siCntl. Pathways selected have FDR < 0.05. Blue indicates pathways related to adipogenesis and lipid metabolism. **e** FPKM values for adipogenic and lipid metabolism related genes detected in both differential gene expression (FC > 1, P < 0.05) and in DAVID analysis (FDR < 0.05) (n = 3/grp). Group comparison was made using One-Way ANOVA where * P < 0.05.

**Figure 5:**
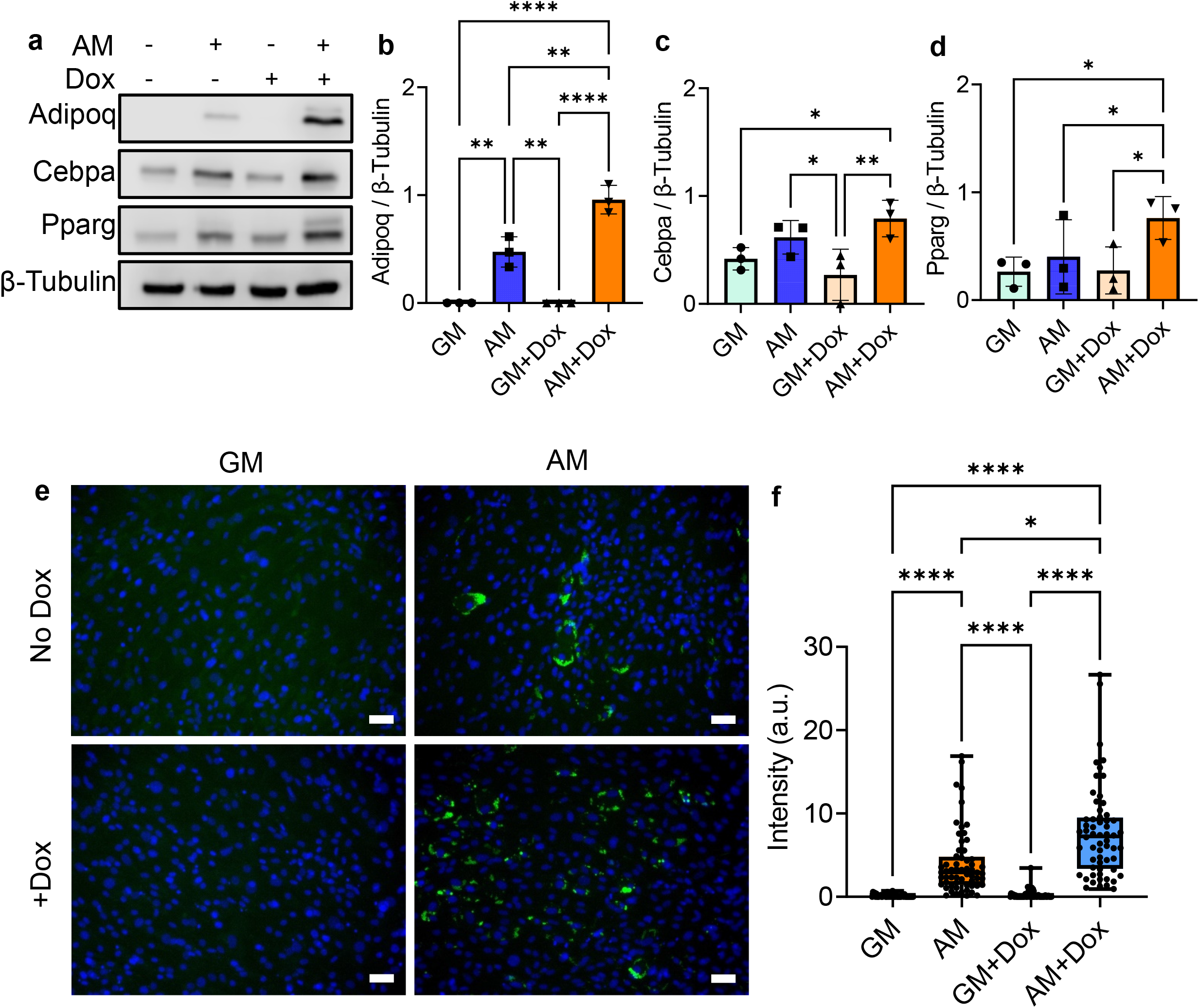
dnKASH expression induces accelerated adipogenesis. **a** Representitive western images of doxycycline induced dnKASH cells and control cells grown in growth media and adipogenic media. **b** During adipogenesis doxycycline treated samples had 98% increased levels of Adipoq (n = 3, P < 0.01). **c** Pparg in doxycycline treatment group during adipogenesis increased by 90% (n = 3, P < 0.05). **d** Cepba experienced an increase of 27% but was not significant during adipogenesis in doxycycline treatment (n = 3, p = 0.38). Representative photos of lipid droplets (green) and DNA (blue). **f** Quantification of mean lipid droplet intensity per cell in each field of view showed an increase of lipid droplet amounts in doxycycline treated cells by 258% (n = 50, P < 0.05) during adipogensis. Western analysis group comparisons were made using One-Way ANOVA. Lipid Droplet Intensity group comparisons were made using Kruskal-Wallis test. * P < 0.05, ** P < 0.01, *** P < 0.001, **** P < 0.0001. Scale bar represents 50μm.

### dnKASH expression accelerates adipogenesis

To quantify the effects of dnKASH expression on adipogenesis, dnKASH-MSCs with ±Dox treatment were compared under AM conditions. Undifferentiated GM groups were also utilized as controls. Shown in **Fig.5a-f**, addition of AM onto no Dox treated MSCs increased adipogenic proteins Adipoq (500%), Cebpa (48%), Pparg (53%) and mean lipid staining per cell by 2400%. With +Dox treatment, AM addition resulted in a larger increases in adipogenic proteins Adipoq (1000%), Cebpa (300%), Pparg (250%) and mean lipid staining per cell by 6000%. Doxycycline alone did not alter the rate of adipogenic differentiation (**Fig. S4**) nor Sun1 and Sun2 levels (**Fig.S5**). Therefore, our results indicate that expression of dnKASH fragments via+Dox treatment resulted in significant increases of Adipoq, Pparg and more than doubled the lipid droplets when compared to no Dox treatment.

To understand the accelerated adipogenesis during dnKASH expression, we performed RNA-seq analysis. DESEQ2 analyses filtered genes pairs with significant expression differentials (fold change ≥ 2-fold, P < 0.05). Shown in **Fig. 6a**, hierarchical heatmap showed a clustering of +Dox treatments (i.e. dnKASH expression) regardless of the media type used. AM cultured cells were further sub-clustered inside their respective ±Dox clades. Shown in **Fig. 6b**, variance of the two principle components were 25.2% and 16.9%. Comparing the gene profiles between AM treated ±Dox groups, DAVID and STRING analyses identified sixteen differentially expressed pathways with an FDR < 0.05 that were either significantly up regulated (**Figs. 6c** **& S6**) and down regulated (**Figs. 6d** **& S7**). +Dox treatment significantly increased the expression of adipogenic genes Adipoq, Fabp4, Lipe, Plin1, Cidec, Agpath2, Acsl1 and mgll genes (**Fig.6e**) and up-regulated lipid metabolism, fatty acid metabolism and PPAR signaling pathways (**Figs. 6c**, blue bars). In order to control for doxycycline effect, expression of Dox-independent dnKASH domain via a secondary plasmid^35^ also showed increased adipogenic gene expression when compared to empty vector (**Fig. S8c**). Showing an opposite that of siSun treatment, the AM +Dox treatment group showed downregulation of inflammatory and immune pathways (**Figs. 6d** **& S6**).Tables of DAVID pathway analysis can be found in **Tables S3** and **S4**.

**Figure 6:**
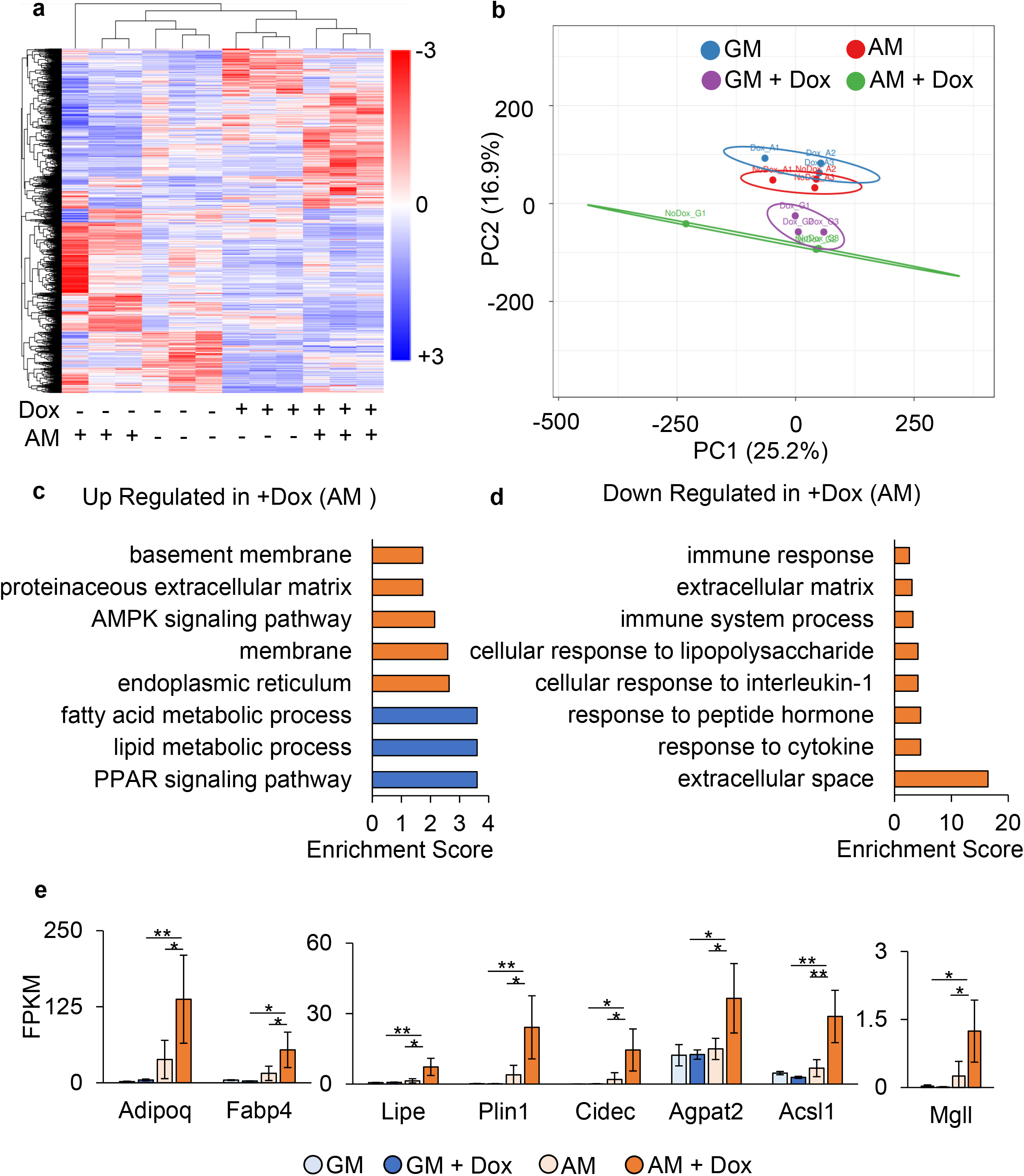
dnkKASH expression increases adipogenesis and lipid metabolism related genes. **a** Heatmap of genes with significant differential (FC > 1 and P < 0.05) gene expression during doxycycline induced dnKASH expression (n = 24). **b** Principle component plot where principal component 1 and principal component 2 explain 25.2% and 16.9% of the total variance, respectively. Prediction ellipses indicate that with a probability of 0.95, a new observation from the same group will fall inside the ellipse. *n* = 24 data points. **c** DAVID analysis of genes up regulated in doxycycline treatment compared to control. Blue indicates pathways related to adipogenesis and lipid metabolism. Pathways selected have FDR < 0.05. **d** DAVID analysis of genes down regulated in doxycycline group compared to control. Pathways selected have FDR < 0.05. **e** FPKM values for adipogenic and lipid metabolism related genes detected in both differential gene expression (FC > 1, P < 0.05) and in DAVID analysis (FDR < 0.05) (n = 3/grp). Group comparison was made using One-Way ANOVA. * P < 0.05, ** P < 0.01.

### H3K9me3 levels and enrichment at the adipogenic gene Adipoq increases during Sun1/2 depletion

To understand possible changes in chromatin during Sun1/2 depletion, we next measured heterochromatin markers H3K9me3, H3K27me3 and the euchromatin marker H3K4me3 in siSun and siCtrl treated MSCs under both GM and AM conditions (**Fig. 7a-d**). Comparing siCntl and siSun counterparts, siSun treatment increased H3K9me3 levels by 56% and 86% in the AM and GM groups. H3K27me3 was decreased by 48% under AM with no changes under GM. No changes were detected in the euchromatin marker H3K4me3.

**Figure 7:**
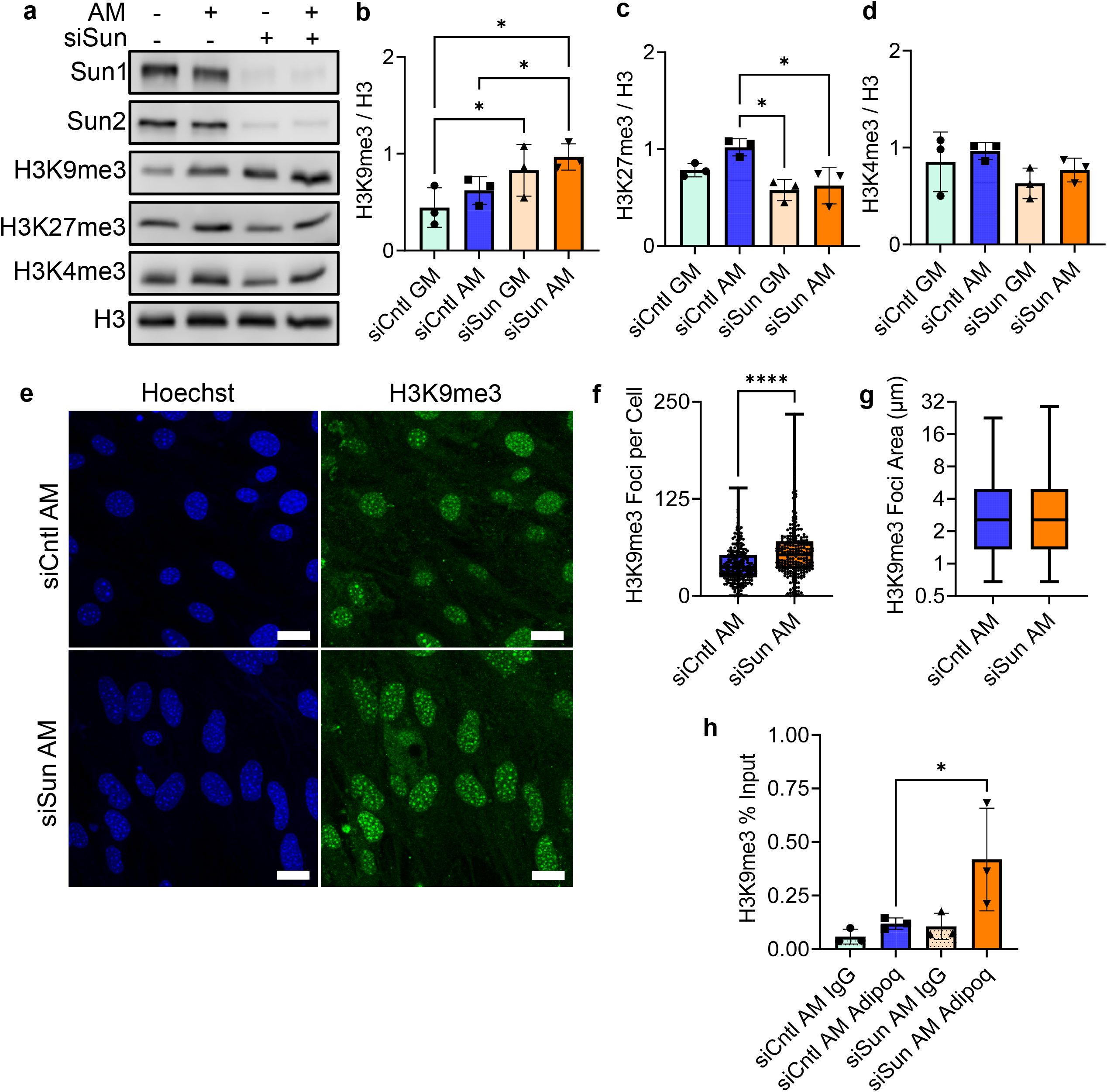
Global levels of H3K9me3 and enrichment on Adipoq increases during Sun1/2 depletion. **a** Representative western images of heterochromatin markers H3K9me3 and H3K27me3 and euchromatin marker H3K4me3 in siSun and siCntl treatments during growth in growth media and adipogenic media. **b** Western analysis of heterochromatin marker H3K9me3 revealed an increase of 56% in siSun cells compared to siCntl during adipogenesis (n = 3, P < 0.05). **c** H3K27me3 had a decrease of 48% in siSun cells compared to siCntl cells during adipogenesis (n =3, P < 0.05). **d** Euchromatin marker H3K4me3 experienced no significant changes in global protein levels between siSun and siCntl-treated cells during adipogenesis. **e** Representative images of siSun and siCntl-treated cells grown in adipogenic media and stained for H3K9me3 (green) and Hoechst 33342 (blue). **f** H3K9me3 foci count per cell in siSun cell compared to Sicntl Cells during adipogenesis increased by 43% (n = 213, P < 0.0001). **g** No detectable increase of H3K9me3 foci area was found in siSun cells during adipogenesis (n = 8460). **h** CUR&RUN-qPCR targeting H3K9me3 localization on Adipoq showed an increase of 156% in siSun cells compared to siCntl (n = 3, P < 0.05). Western analysis group comparisons were made using One-Way ANOVA. H3K9me3 Foci count and area comparisons were made using Mann-Whitney Test. CUR&RUN-qPCR comparisons were done using One-Tailed Students T-Test. * P < 0.05, ** P < 0.01, *** P < 0.001, **** P < 0.0001. Scale bar represents 25μm.

To further investigate the increased H3K9me3 levels in the siSun group, we quantified confocal images of H3K9me3 (green) and DNA (blue) under AM conditions (**Fig. 7e-g**). siSun treatment increased H3K9me3 foci count by 43%, foci area was unchanged. To detect if increased H3K9me3 led to increased enrichment on adipogenic genes, we probed the most increased gene under AM, Adipoq, via a CUT&RUN extraction targeting H3K9me3. Shown in **Fig. 7k**, H3K9me3 enrichment on Adipoq was increased by 156% in the siSun AM group compared to the siCntl AM group.

### H3K9me3 Levels and Enrichment at the adipogenic gene Adipoq remains unaltered during dnKASH disruption of the LINC Complex

Investigating the increased adipogenesis of dnKASH-MSCs, heterochromatin markers H3K9me3, H3K27me3 and euchromatin marker H3K4me3 were measured under both GM and AM conditions (**Fig. 8a-d**). Under GM conditions no changes were detected. In contrast to Sun1/2 depletion, +Dox treatment decreased H3K9me3 by 51% under AM. AM+Dox treatment also decreased H3K27me3 by 56%. No H3K4me3 differences were detected.

**Figure 8:**
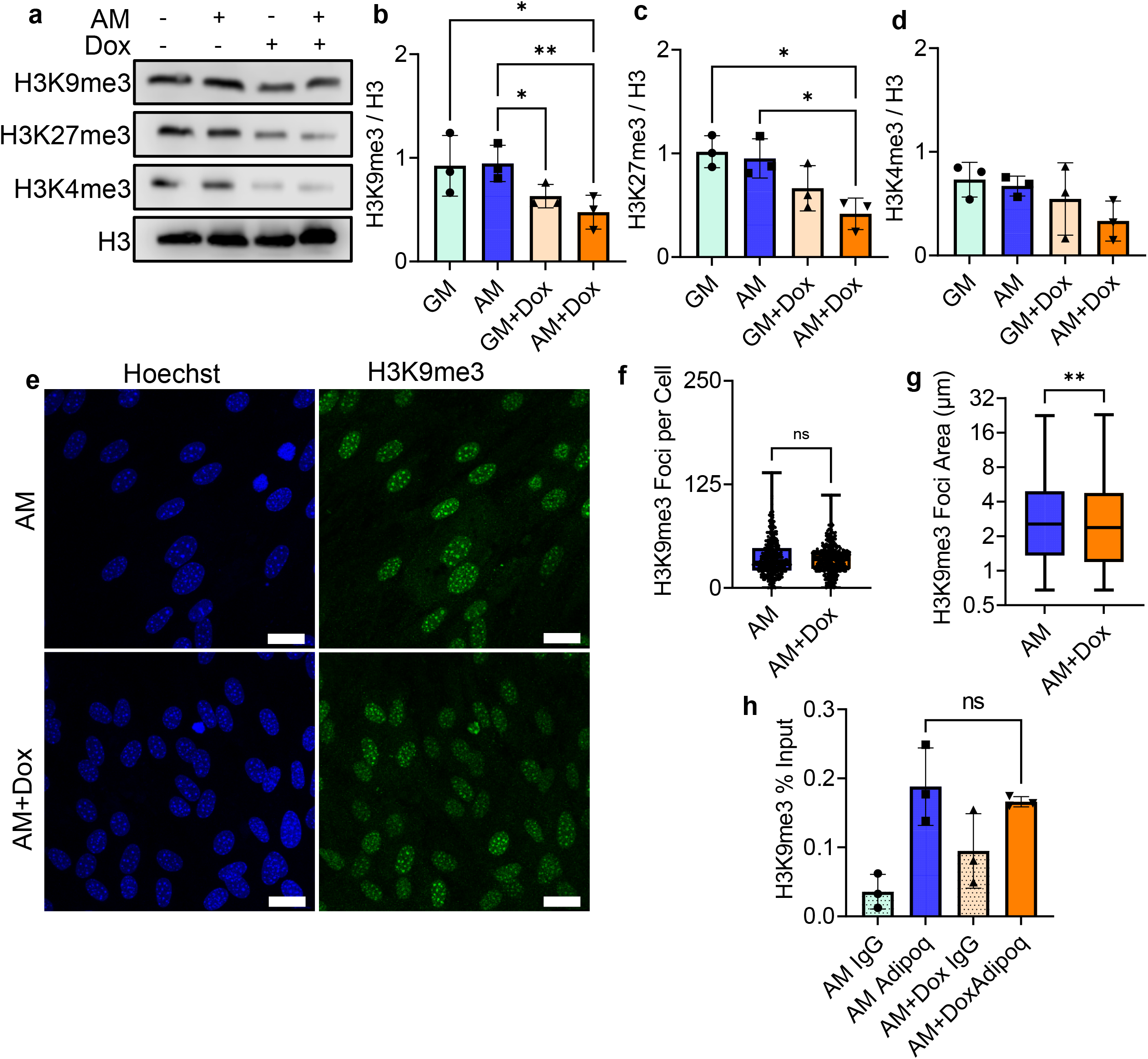
H3K9me3 Levels are unaltered during dnKASH expression. **a** Representative images of doxycycline-induced dnKASH expression of heterochromatin markers H3K9me3 and H3K27me3 and euchromatin marker. **b** H3K9me3 levels decreased during adipogenesis in the doxycycline treatment group compared to controls by 51% (n =3, P < 0.01). **c** H3K27me3 levels decreased by 56% in the doxycycline treatment group compared to control during adipogenesis (n = 3, P < 0.05). **d** H3K4me3 levels had no significant changes in doxycycline. **e** Representative images of doxycycline-treated cells and controls cells stained for H3K9me3 (green) and Hoescht (blue) during growth in adipogenic media. **f** H3K9me3 foci count per cell did not show significant changes between the doxycycline treatment group and the control group during growth in adipogenic media (n = 328). **g** The doxycycline treatment group had decreased H3k9me3 foci area during adipogenesis by 7% compared to control (n = 11317, P < 0.01). **h** CUR&RUN-qPCR targeting H3K9me3 localization on Adipoq showed no significant increase in doxycycline treated cells compared to controls (n = 3). Western analysis group comparisons were made using One-Way ANOVA. H3K9me3 Foci count and area comparisons were made using Mann-Whitney Test. CUR&RUN-qPCR comparisons were done using One-Tailed Students T-Test. * P < 0.05, ** P < 0.01, *** P < 0.001, **** P < 0.0001. Scale bar represents 25μm.

Confocal imaging of H3K9me3 (green) and DNA (Blue) in AM (**Fig. 8e-g**) revealed no H3K9me3 foci count changes, foci area was increased by 7%. Further, Methylation of H3K9 associated with Adipoq was not different when dnKASH was expressed during adipogenic constraint. (**Fig. 8h**).

## Discussion

Both Sun and Nesprin – connected to Sun via its KASH domain – are part of the LINC complex function. Depletion of Sun proteins and expression of dominant negative KASH isoforms (i.e. dnKASH) both interfere with transmittance of mechanical information to cell nucleus. In addition to their function in nucleo-cytoskeletal connectivity, Sun proteins have specific connections to chromatin and may participate in gene availability. Here we report that depletion of Sun1/2 results in increased H3K9me3 accrual at both global and gene scales. dnKASH treatment has no such effect, suggesting that these two methods cannot be used interchangeably in studying LINC function.

Observed increase in nuclear area, perimeter and decreased circularity seen during Sun1/2 depletion is similar to nuclear morphology changes during Lamin A/C depletion^58,59^. Previous research has shown that both depletion of Lamin A/C and disassociation of Sun1 from nuclear envelope via Sun1-KDEL expression softens nuclei^49^, suggesting that Sun proteins – Like LaminA/C – supports nuclear structure and possibly tether chromatin. We have found that disabling LINC function via dnKASH decreases nuclear area, consistent with loss of F-actin tension^60^ and increased nuclear height^61,62^, but result in smaller changes in perimeter and circularity suggesting a structurally intact nuclei. Prior work utilizing micropipette-induced nuclear deformations supports the idea that dnKASH expression does not significantly reduce nuclear stiffness^49^. Indeed, we have previously shown that depletion of Sun1/2 causes a reduction in isolated nuclear stiffness while dnKASH expression does not ^48^. Thus, our data indicates dnKASH regulates nuclear morphology and adipogenesis without reducing nuclear stiffness while depletion of Sun1/2 is potentially able to regulate adipogenesis and nuclear shape by softening nuclei.

Previous studies have shown that depletion of Sun1/2 reduces tethering of chromosomes to the nuclear envelope^53,63,64^, alters nucleolus morphology^65^, and increases H3K9me3 levels in hTERT-RPE1 and MCF10A cells^66^. Our results show similar trends where Sun1/2 depletion increased both H3K9me3 protein levels and number of H3K9me3 heterochromatic foci. In contrast, no H3K9me3 changes were observed under dnKASH expression.

Our data indicates that MSCs from bone marrow are primed for adipogenic transformation under adipogenic media conditions. Disrupting nucleo-cytoskeletal connectivity via dnKASH accelerates this process. dnKASH expression was not accompanied with large changes in chromatin. As dnKASH expression limits the nuclear entry of β-Catenin^33^ and Yap^67^ with anti-adipogenic functions in MSC^29–32^, disabling LINC function may accelerate adipogenesis through these mechanisms. In contrast, increased H3K9me3 methylation under Sun1/2 depletion impeded adipogenic differentiation as shown through western blot analysis, RNA-seq analysis, and lipid droplet counts. Consistently, we observed H3K9me3 heterochromatin enrichment on the adipogenic gene *Adipoq*, presumably decreasing availability for adipogenic transformation. These results indicate that Sun1/2 proteins regulate H3K9me3 heterochromatin organization independent of the LINC-mediated nucleo-cytoskeletal complex connectivity and alter adipogenesis in MSCs.

Downregulation of LINC complex components including Sun1, Sun2 and Nesprin2 have been observed in early stage human carcinomas^68,69^ and loss of these proteins were associated with more aggressive tumors and poorer prognosis^70–73^. In our study, we found that Sun1/2 depletion increased tumorigenic and osteolytic factors *Cxcl10*, *Cxcl1*, and *Cxcl5* (Table S1). Increases in these genes are associated with increased metastasis in breast cancer^74^ and prostate cancer^75^ as well as promoting osteoclast differentiation^76,77^ and angiogenesis^78^. In contrast, dnKASH expression downregulated the same genes (Table S4) indicating a reduction in the osteolytic and tumorigenic signaling. While the underlying mechanism for these observed difference between Sun1/2 and dnKASH treatments are unknown, our results indicate that Sun proteins may play a more direct function in activating cancer-associated signaling in MSCs.

In conclusion, this work reveals new insight into the role of Sun1/2 proteins in the nuclear interior and differentiation. Depletion of Sun1/2 increased H3K9me3 heterochromatin and inhibited adipogenesis. Our findings show that these outcomes were different in dnKASH expressing MSCs and thus the effects of Sun1/2 depletion may be independent of the LINC complex function. These results have potential impacts on our understanding enteropathies targeting the nuclear envelope, such as heightened symptoms seen in Emery-Dreifuss muscular dystrophy (EDMD) patients with additional Sun mutations or ^64,79^ aberrant accumulation of Sun proteins at the nuclear envelope associated with more severe progeria symptoms^80,81^. Thus, these results expand our understandings of the important role inner nuclear membrane proteins have in regulating proper nuclear functions and ultimately human health.

## Materials and Methods

### MSCs Isolation

Bone marrow derived MSCs (mdMSC) from 8-10 wk male C57BL/6 mice were isolated as described^58,82^ from multiple mouse donors and MSCs pooled, providing a heterogenous MSCs cell line. Briefly, tibial and femoral marrow were collected in RPMI-1640, 9% FBS, 9% HS, 100 μg/ml pen/strep and 12μM L-glutamine. After 24 hours, non-adherent cells were removed by washing with phosphate-buffered saline and adherent cells cultured for 4 weeks. Passage 1 cells were collected after incubation with 0.25% trypsin/1 mM EDTA × 2 minutes, and re-plated in a single 175-cm2 flask. After 1-2 weeks, passage 2 cells were re-plated at 50 cells/cm2 in expansion medium (Iscove modified Dulbecco’s, 9% FBS, 9% HS, antibiotics, L-glutamine). mdMSC were re-plated every 1-2 weeks for two consecutive passages up to passage 5 and tested for osteogenic and adipogenic potential, and subsequently frozen.

### Stable dnKASH Cell Line

MSCs were stably transduced with a doxycycline inducible plasmid expressing an mCherry tagged dominant-negative KASH domain. dnKASH plasmid was lentiviral packaged as a generous gift from Dr. Daniel Conway. Vector map found here: https://www.addgene.org/125554/. Lentivius supernatant was added to growth media with polybrene (5 μg/ml). Lentivirus growth media mixture was added to 50-70% confluent MSCs. Lentivirus media was replaced 48 hours later with selection media containing G418 (1mg/ml) for 5 days to select stably infected dnKASH-MSCs.

### Cell Culture, Pharmacological Reagents, and Antibodies

Fetal calf serum (FCS) was obtained from Atlanta Biologicals (Atlanta, GA). MSCs were maintained in IMDM with FBS (10%, v/v) and penicillin/streptomycin (100μg/ml). For immunostaining experiments, seeding cell density was 3,000 cells per square centimeter. For adipogenic differentiation experiments, the seeding cell density was 10,000 cells per square centimeter. Cells were either grown in growth media (GM) or adipogenic media (AM) Cells were transfected 24 hours after cell seeding with siRNA targeting Sun1 and Sun2 (siSun) or a control sequence (siCntl) using RNAiMax from Invitrogen. Adipogenic media was placed on siRNA treated cells twenty four hours after the transfection, the adipogenic media was added which contained dexamethasone (0.1μM), insulin (5 μg/ml), and indomethacin (1 μg/ml) for 5 days.

For dnKASH-MSCs, cells were seeded at 10,000 cells per square centimeter. Twenty four hours after seeding, dnKASH cells were given growth media containing doxycycline (1 μg/ml). Adipogenic media containing dexamethasone (0.1μM), insulin (5 μg/ml), indomethacin (1 μg/ml), and doxycycline (1 μg/ml) (AM+Dox) or growth media (GM+Dox) was placed on dnKASH-MSCs twenty four hours after adding initial doxycycline. Control cells were grown in growth media (GM) or adipogenic media (AM) without doxycycline. Growth media or adipogenic media with or without fresh doxycycline were changed every 48 hours.

### siRNA Silencing Sequences

For transient silencing of MSCs, cells were transfected with gene-specific small interfering RNA (siRNA) or control siRNA (20 nM) using RNAiMax (ThermoFisher) according to manufacturer’s instructions. The following Stealth Select siRNAs (Invitrogen) were used in this study: SUN1 (ThermoFischer Scientific, Assay ID #s94913), SUN2 (ThermoFischer Scientific, Assay ID s104591).

### qPCR

2ul of each CUT&RUN sample was run in 20ul reaction following Bio-Rad protocols targeting Adipoq (Bio-Rad, 10025636). Briefly, 20ul reactions were made using SsoAdvanced Master Mix (Bio-Rad, 1725270). Reactions were then run at 95°C for two minutes. Then samples were heated at 95°C for 15 seconds then cooled to 60°C for 30 seconds which both steps were repeated for 40 cycles. Finally, samples were run at 60°C for two minutes. Samples were then analyzed for percent of input sample for CUT&RUN-qPCR.

### RNA-seq

Five days after adipogenic treatment, following the above protocols, total RNA was extracted using RNAeasy (Qiagen) for three samples per group. Total RNA samples were sent to Novogene for mRNA sequencing and analysis. Briefly, index of the reference genome was built using Hisat2 v2.0.5 and paired-end clean 2 reads were aligned to the reference genome using Hisat2 v2.0.5. featureCounts v1.5.0-p3 was used to count the reads numbers mapped to each gene. And then FPKM of each gene was calculated based on the length of the gene and reads count mapped to this gene. FPKM, expected number of Fragments Per Kilobase of transcript sequence per Millions base pairs sequenced. Differential expression analysis was performed using the DESeq2 R package (1.20.0). DESeq2 provides statistical routines for determining differential expression in digital gene expression data using a model based on the negative binomial distribution. The resulting P-values were adjusted using the Benjamini and Hochberg’s approach for controlling the false discovery rate. Genes with an adjusted P-value < = 0.05 found by DESeq2 were assigned as differentially expressed. Genes with significant differential gene expression were further analyzed with DAVID^83^ for pathway analysis. Pathways with a FDR < = 0.05 were selected.

### Immunofluorescence

Twenty four hours after the siRNA treatment against Sun1/Sun2 or dnKASH expression, cells were fixed with 4% paraformaldehyde. Cells were permeabilized by incubation with 0.3% Triton X-100. Cells were incubated in a blocking serum in PBS with 5% Donkey Serum (017-000-121, Jackson Immuno Research Laboratories). Primary antibody solution was incubated on the cells for 1h at 37oC, followed by secondary antibody incubation of either Alexa Flour 594 goat anti-rabbit (Invitrogen), Alexa Fluor 488 goat anti-mouse (Invitrogen), Alexa Fluor 488 chicken anti-rabbit (Invitrogen), or Alexa fluor 594 Donkey anti-mouse (Invitrogen). For nuclear staining cells were incubated with NucBlue Hoechst 33342 stain (Fisher Scientific). Primary and secondary concentrations were both 1:300 by volume.

### Image Analysis

Five days after adding adipogenic media, cells were fixed and stained with Lipid Spot 488 (Biotium, #70069), and NucBlue Hoechst 33342 stain. Images were taken using 20x objective and were exported to quantify lipid droplet formation via a custom-made MATLAB program (The MathWorks, Natick, MA), previously published^35,58^. The minimum pixel intensity of 80 was used to isolate lipid droplet staining. The mean lipid droplet intensity per cell was calculated by dividing the sum of lipid droplet stain intensity by the nuclei count per image. Exported images were used to quantify lipid droplet formation, Sun1, Sun2, mCherry, nuclear area, nuclear perimeter, and nuclear circularity via the custom-made MATLAB program previously published^58^. Cell Profiler (https://cellprofiler.org/) was used to count the number of H3K9me3 foci per cell and foci area.

### Western Blotting

Whole cell lysates were prepared using radio immunoprecipitation assay (RIPA) lysis buffer (150mM NaCl, 50mM Tris HCl, 1mM EDTA, 0.24% sodium deoxycholate,1% Igepal, pH 7.5) to protect the samples from protein degradation NaF (25mM), Na3VO4 (2mM), aprotinin, leupeptin, pepstatin, and phenylmethylsulfonylfluoride (PMSF) were added to the lysis buffer. Western protein amounts were normalized to 15μg through BCA Protein Assay (Thermo Scientific, #23225). Whole cell lysates (20μg) were separated on 10% poly-acrylamide gels and transferred to polyvinylidene difluoride (PVDF) membranes. Membranes were blocked with milk (5%, w/v) diluted in Tris-buffered saline containing Tween20 (TBS-T, 0.05%). Blots were then incubated overnight at 4°C with appropriate primary antibodies. Following primary antibody incubation, blots were washed and incubated with horseradish peroxidase-conjugated secondary antibody diluted at 1: 5,000 (Cell Signaling) at RT for 1h in 5% milk in TBST-T. Chemiluminescence was detected with ECL plus (Amersham Biosciences, Piscataway, NJ). At least three separate experiments were used for densitometry analyses of western blots and densitometry was performed via NIH ImageJ software.

### CUT&RUN

CUT&RUN was performed using the CUT&RUN Assay Kit (Cell Signaling #86652). Briefly, cells were harvested and centrifuged, washed, and bound to Concanavalin A-coated magnetic beads. Cells were then permeabilized with digitonin and incubated with primary antibody at 4°C for two hours. Cells were then washed and resuspended with pAG-MNase enzyme and incubated at 4°C for one hour. Cells were then incubated at 37°C for 10 minutes to elute DNA into solution. Solution was then extracted and purified using DNA Purification Buffers and Spin Columns (ChIP, CUT&RUN, Cell Signaling #14209). DNA samples were then used for qPCR or sequencing.

### Statistical Analysis and Reproducibility

Results for densitometry were presented as mean ± SEM. Densitometry and other analyses were performed on at least three separate experiments. Differences between groups were identified by One-Way ANOVA. Analysis of nuclear morphology histone modifications were done using Whitney-Mann test and results were presented as mean ± STD. Differential gene expression analysis via DESEQ2 was done using Wald test. P-values of less than 0.05 were considered significant. Lipid image analysis groups were analyzed via the Kruskal-Wallis test. CUT&RUN-qPCR was analyzed via One-Tailed Students T-Test. All experiments were conducted in triplicate to assure reproducibility.

## Data Availability

RNA-Seq data that support the findings of this study have been deposited in GEO with the accession codes GSE193505.

## Acknowledgements

This study was supported by AG059923, AR075803, P20GM109095, NSF1929188 and, NSF 2025505

## Contributions

Matthew Goelzer: concept/design, data analysis/interpretation, manuscript writing

Sean Howard: data analysis, final approval of manuscript

Anamaria Zavala: data analysis/interpretation, final approval of manuscript

Daniel Conway: concept/design, final approval of manuscript

Janet Rubin: concept/design, data analysis/interpretation, final approval of manuscript

Andre J van Wijnen: concept/design, data analysis/interpretation, final approval of manuscript

Gunes Uzer: concept/design, data analysis/interpretation, financial support, manuscript writing, final approval of manuscript

## Supplementary Figures

**Fig. S1.**
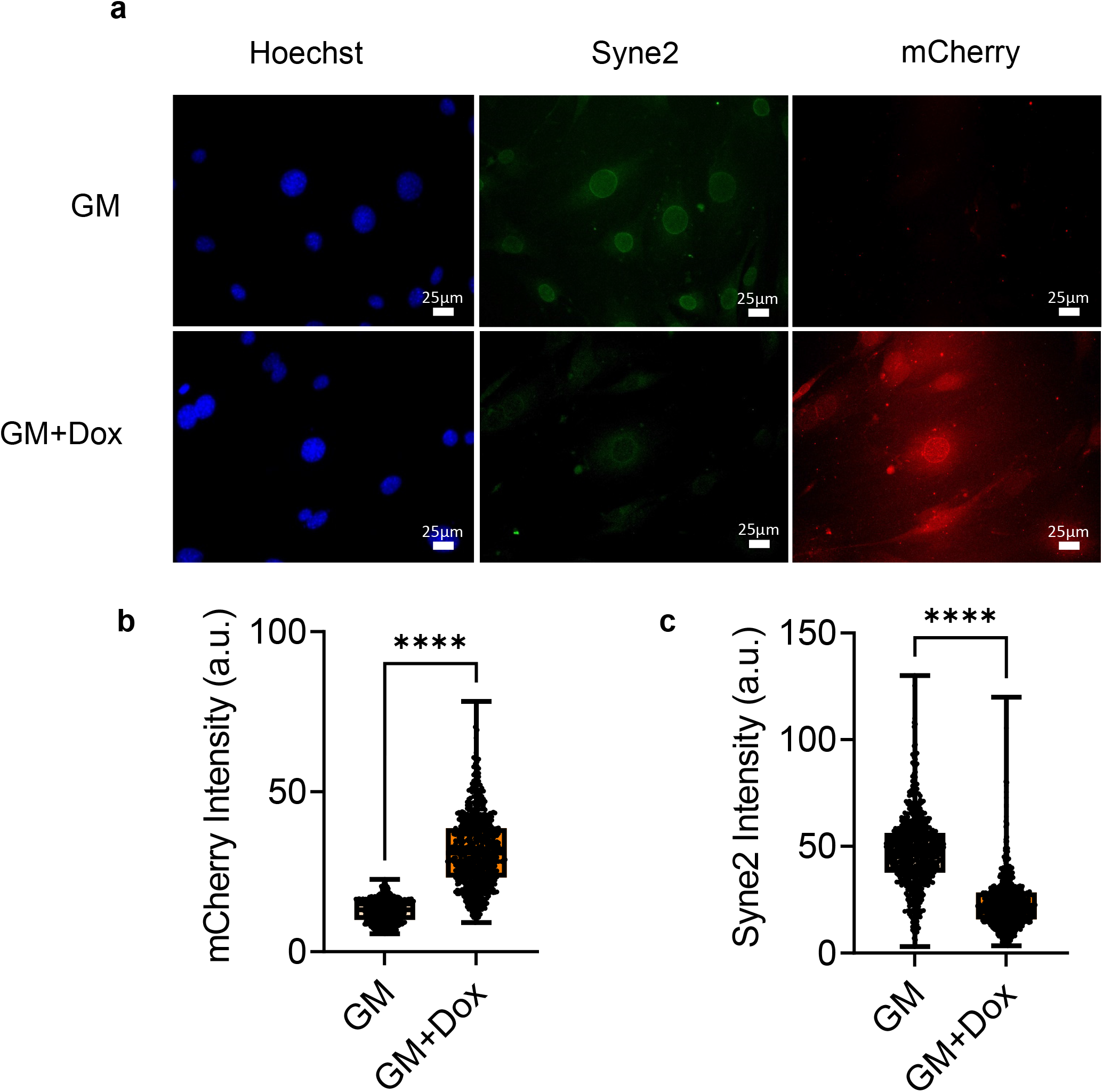
dnKASH expression displaces nesprin2 (Syne2) from nuclear envelope. **a** Representative confocal images of nuclei (Hoechst), Syne2 (Green), and dnKASH (mCherry) in GM and GM+Dox treated MSCs. **b** mCherry intensity per nucleus. GM+Dox MSCs had 200% increase of mCherry intensity (n = 1400, P < 0.0001). **c** Nesprin2 (Syne2) inteisty levels per nucleus. GM+Dox cells had 62% decrease in intensity compared to GM group (n – 1400, P < 0.0001). Groups were compared using Two-Tailed Students T-Test where **** = P < 0.0001.

**Fig. S2.**
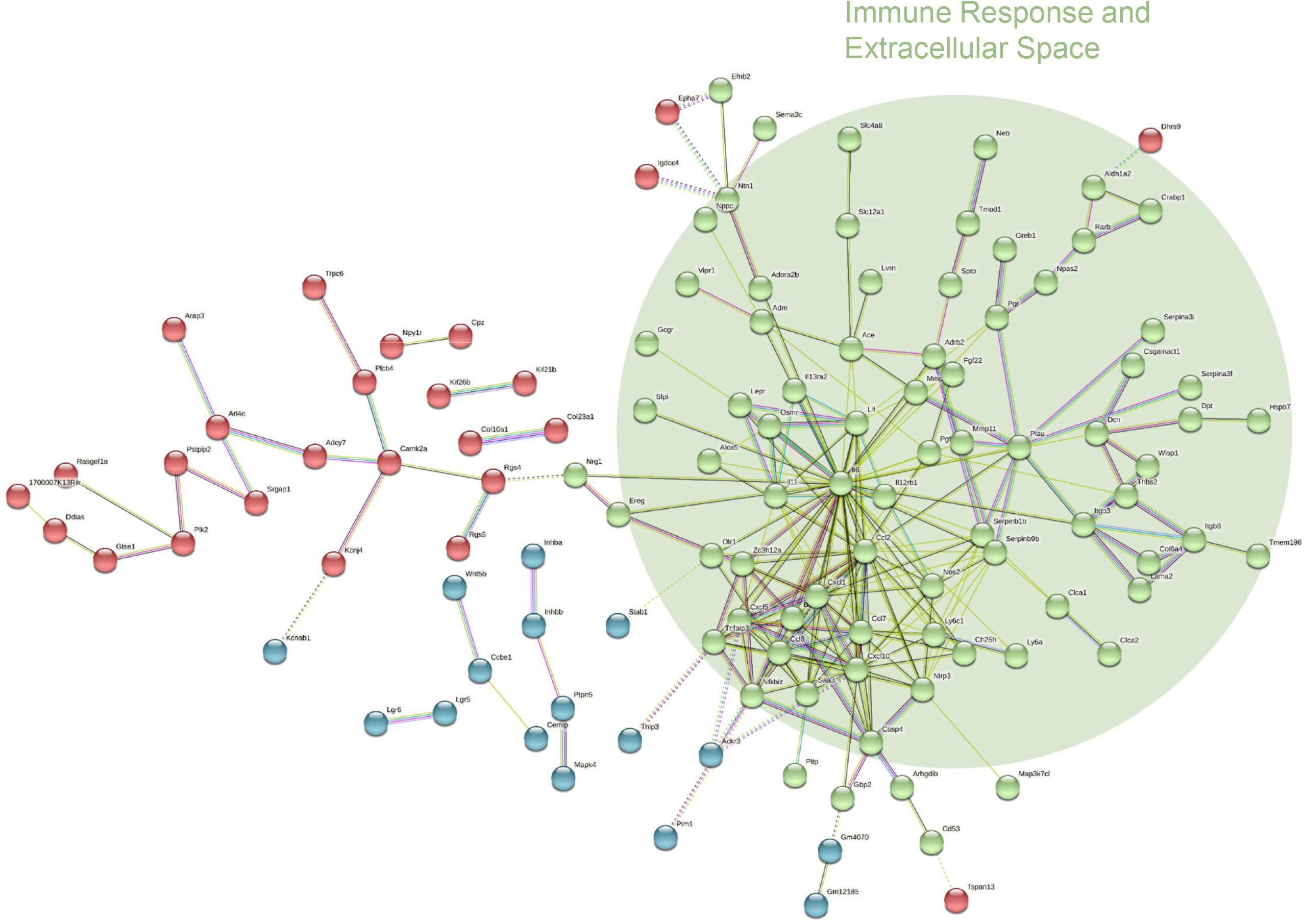
STRING analysis of up regulated genes in siSun AM vs siCntl AM. Green cluster of genes were associated with immune system response and extracellular space (FDR < 0.05).

**Fig. S3.**
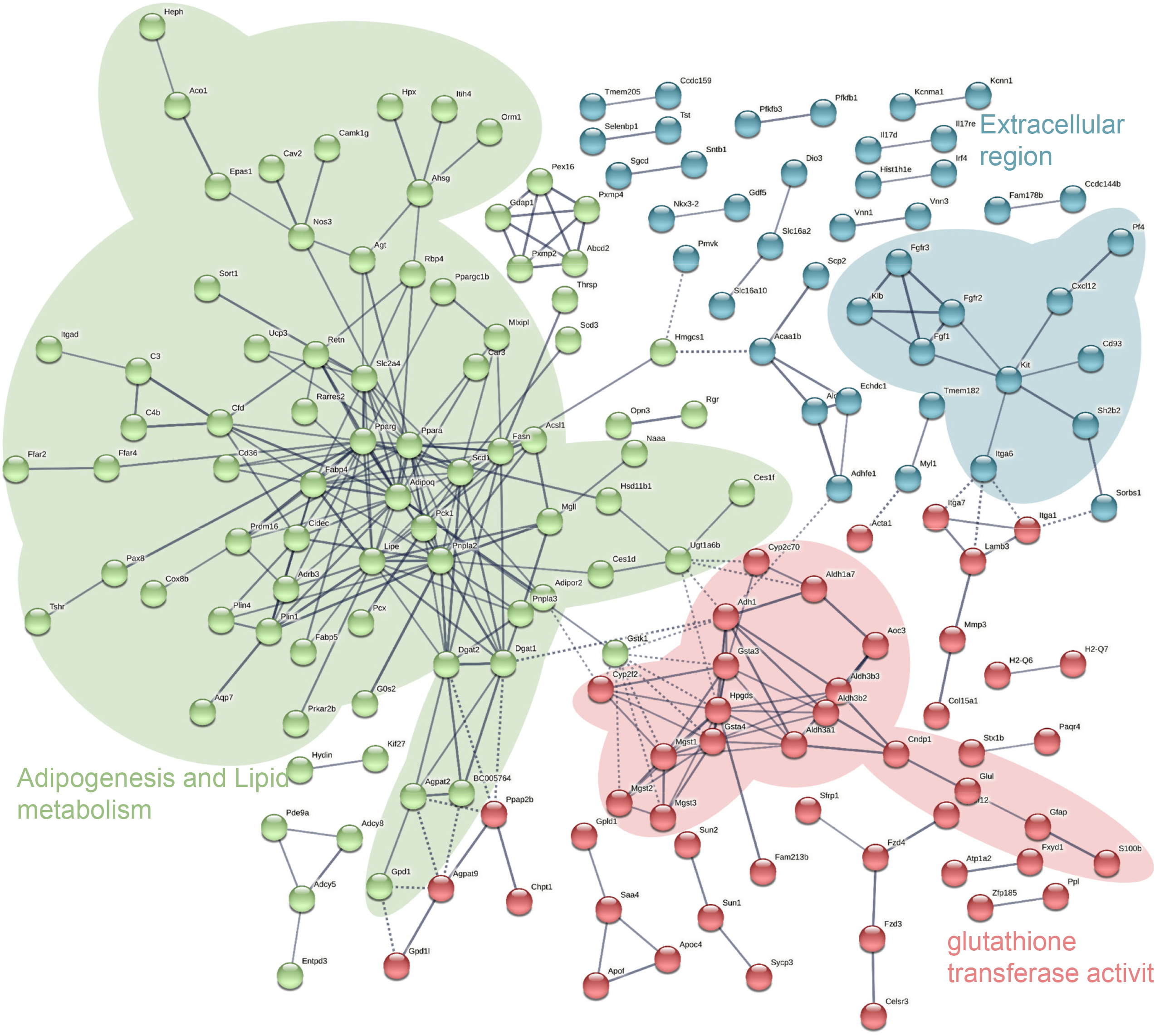
STRING analysis of down regulated genes in siSun AM vs siCntl AM. Green cluster of genes were associated with adipogenesis and lipid metabolism pathways. Blue cluster represents genes associated with extracellular region. Red cluster identifies genes associated with glutathione transferase activity pathway (FDR < 0.05).

**Fig. S4.**
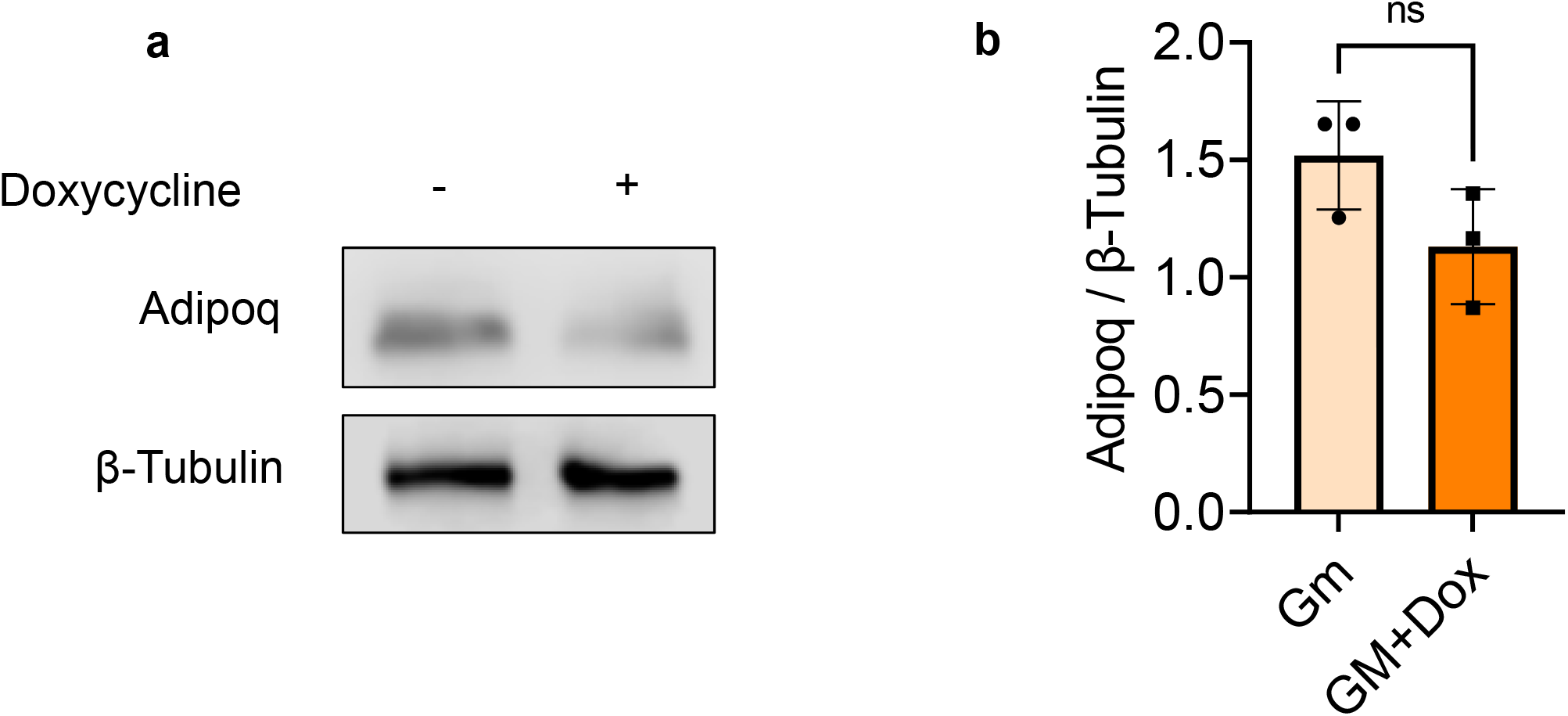
Determination of doxycyline treatment on adipogenesis in MSCs. **a** Representative western blot images of Adipoq during growth in adipogenic media with our without doxycycline. **b** Western blot analysis of Adipoq protein levels during in MSCs grown in adipogenic media with and without doxycycline. No significant differences were detected between AM+Dox or AM groups. Groups were compared using Two-Tailed Students T-Test. * P < 0.05, ** P < 0.01, *** P < 0.001, **** P < 0.0001.

**Fig. S5.**
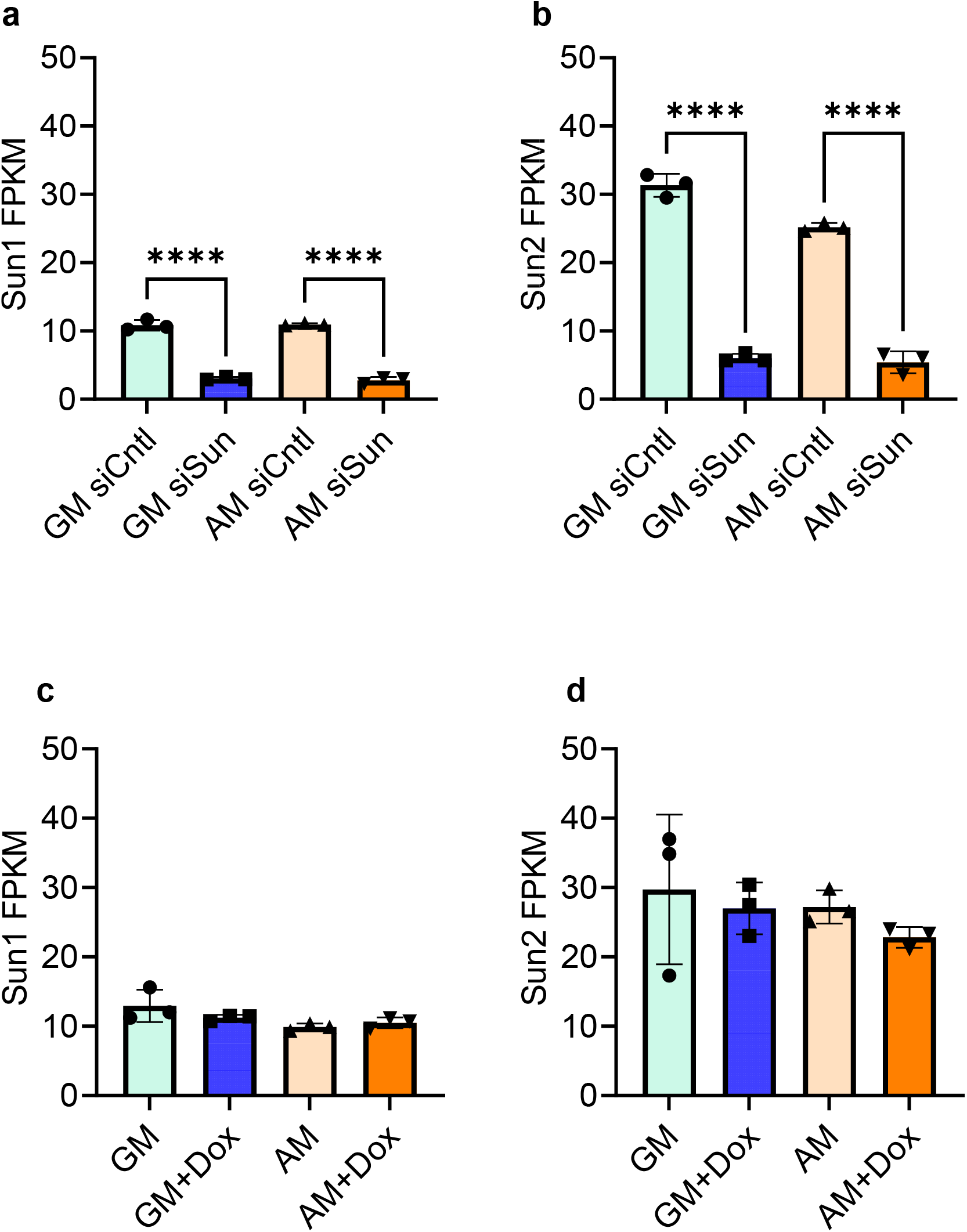
Determination of Doxycyline Treatment on Sun levels. Treating MSCs with siSun decreased both **a** Sun1 and **b** Sun2 gene expression levels. +Dox treatment had no effect on **c** Sun1 and **d** Sun2 gene expression levels. Group comparisons were performed via One-Way ANOVA. * P < 0.05, ** P < 0.01, *** P < 0.001, **** P < 0.0001.

**Fig. S6.**
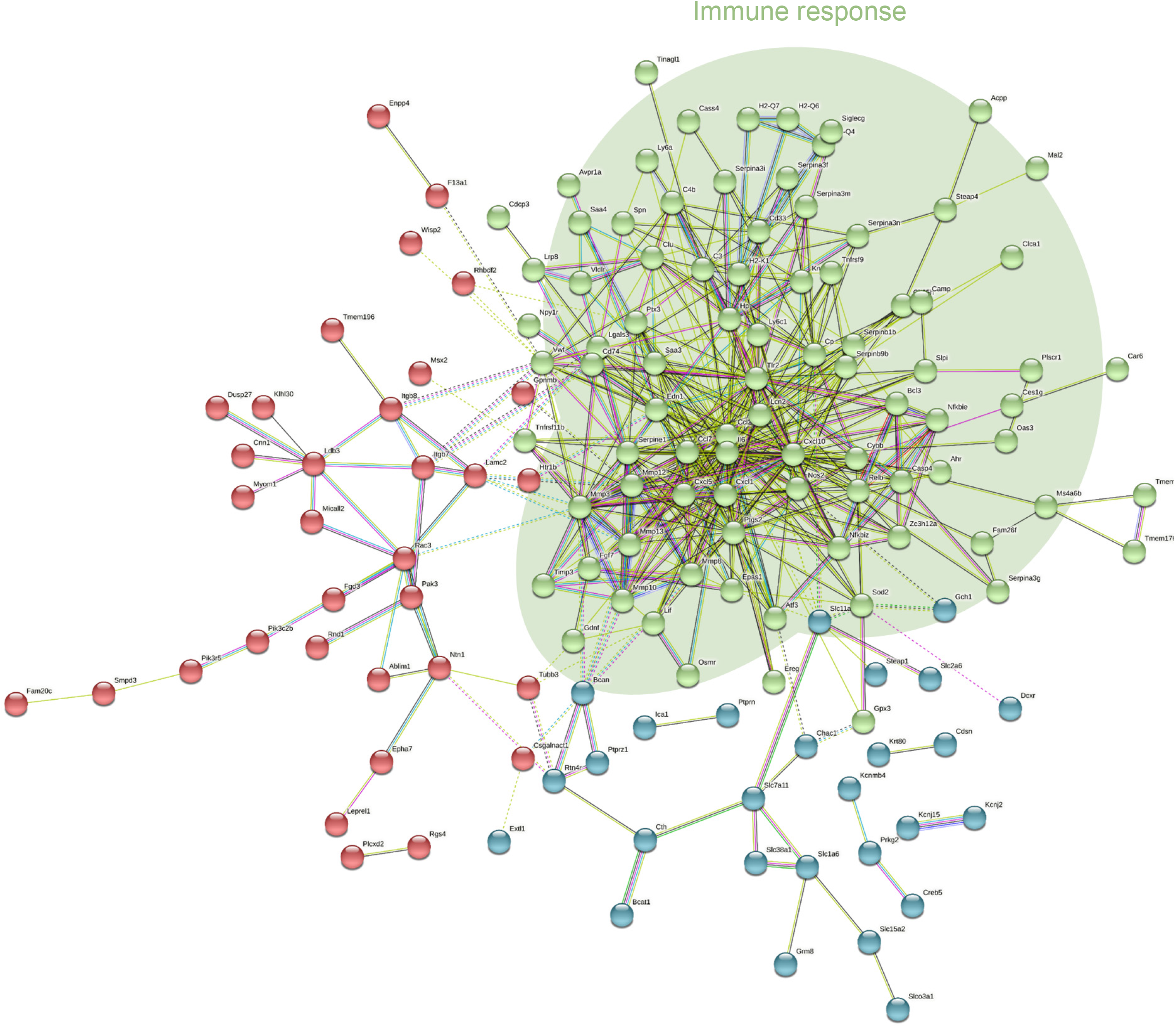
STRING analysis of down regulated genes in AM+Dox vs AM. Green cluster represents genes identified in immune response pathway (FDR < 0.05).

**Fig. S7.**
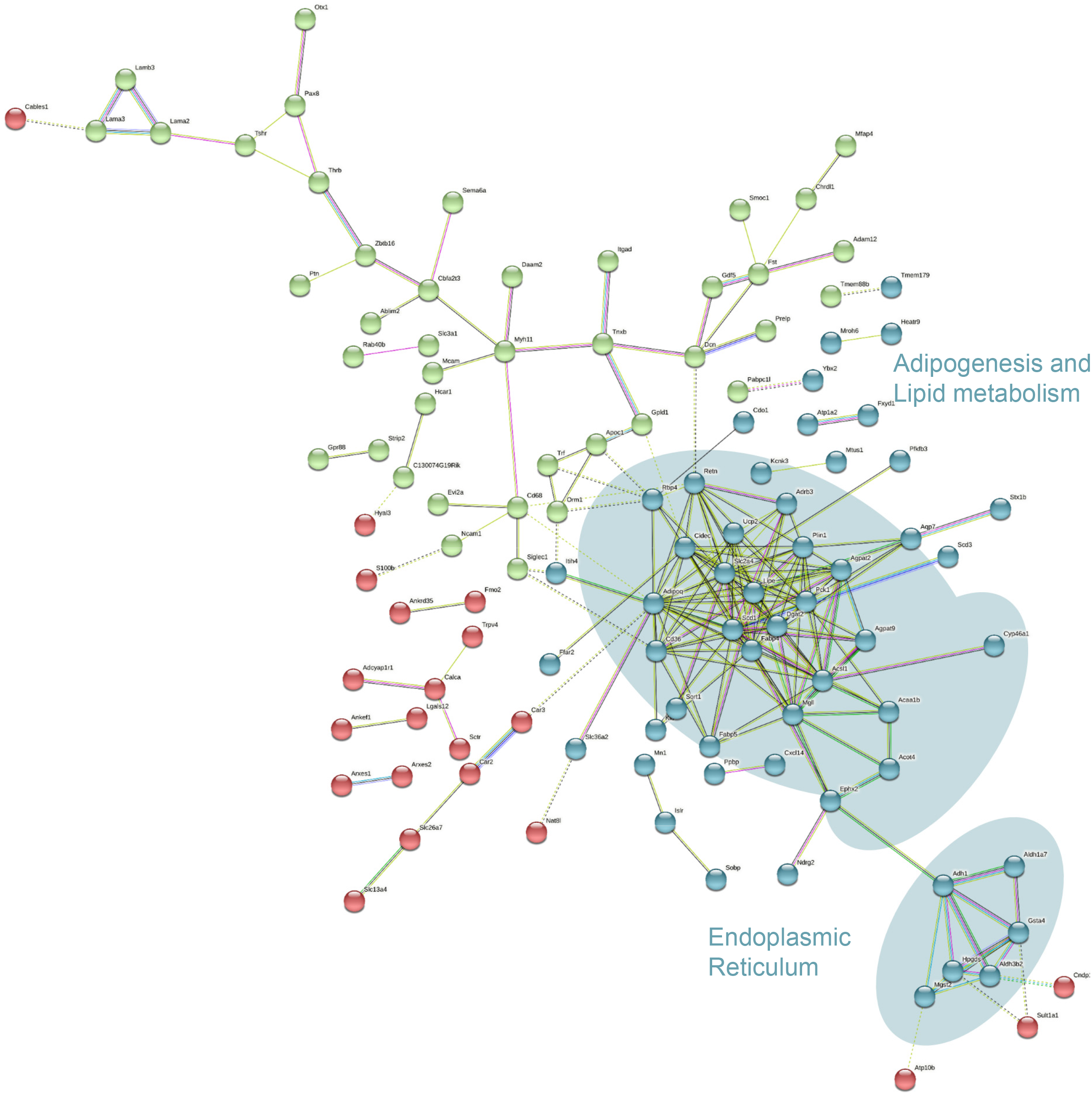
STRING analysis of up regulated genes in AM+Dox vs AM. Blue cluster of genes were associated with adipogenesis and lipid metabolism pathways and endoplasmic reticulum (FDR < 0.05).

**Fig. S8.**
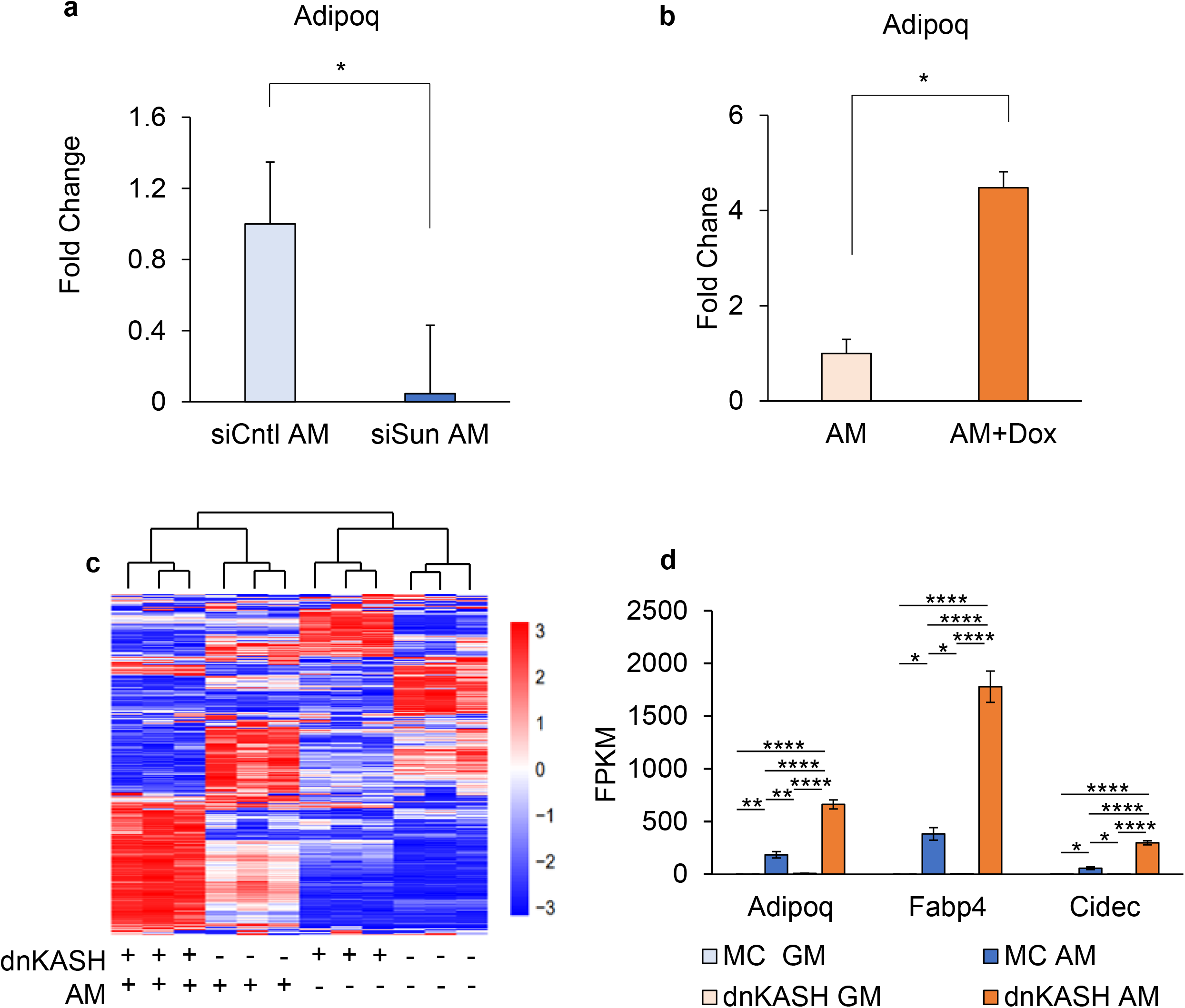
Validation of siSun and dnKASH adipogenic gene expression. **a** qPCR validation of *Adipoq* levels during growth in AM and siSun treatment. *Adipoq* expression decreased by 90% (n = 3, P < 0.05). **b** qPCR validation of Adipoq levels during growth in AM or AM+Dox media. *Adipoq* expression in dnKASH expressing MSCs had 4-fold increase of Adipoq expression (n = 3, P < 0.05). **c** Heatmap of MSCs transfected with secondary plasmid expressing dnKASH or empty vector (MC) during growth in GM or AM media. Genes selected had FPKM > 0.3 and P < 0.05 compared to controls. **d** FPKM levels of *Adipoq, Fabp4,* and *Cidec* during transfection with either dnKASH or MC in GM or AM media. *Adipoq, Fabp4,* and *Cidec* experienced increases of 280% (n =3, P < 0.0001), 400% (n =3, P < 0.0001), and 500% (n =3, P < 0.0001), respectively, in FPKM levels between MC AM and dnKASH AM groups. qPCR validation of siSun and dnKASH Groups were compared using Two-Tailed Students T-Test. FPKM analysis of adipogenic genes *Adipoq, Fabp4,* and *Cidec* was done by One-Way ANOVA. * P < 0.05, ** P < 0.01, *** P < 0.001, **** P < 0.0001.

**Fig. S9.**
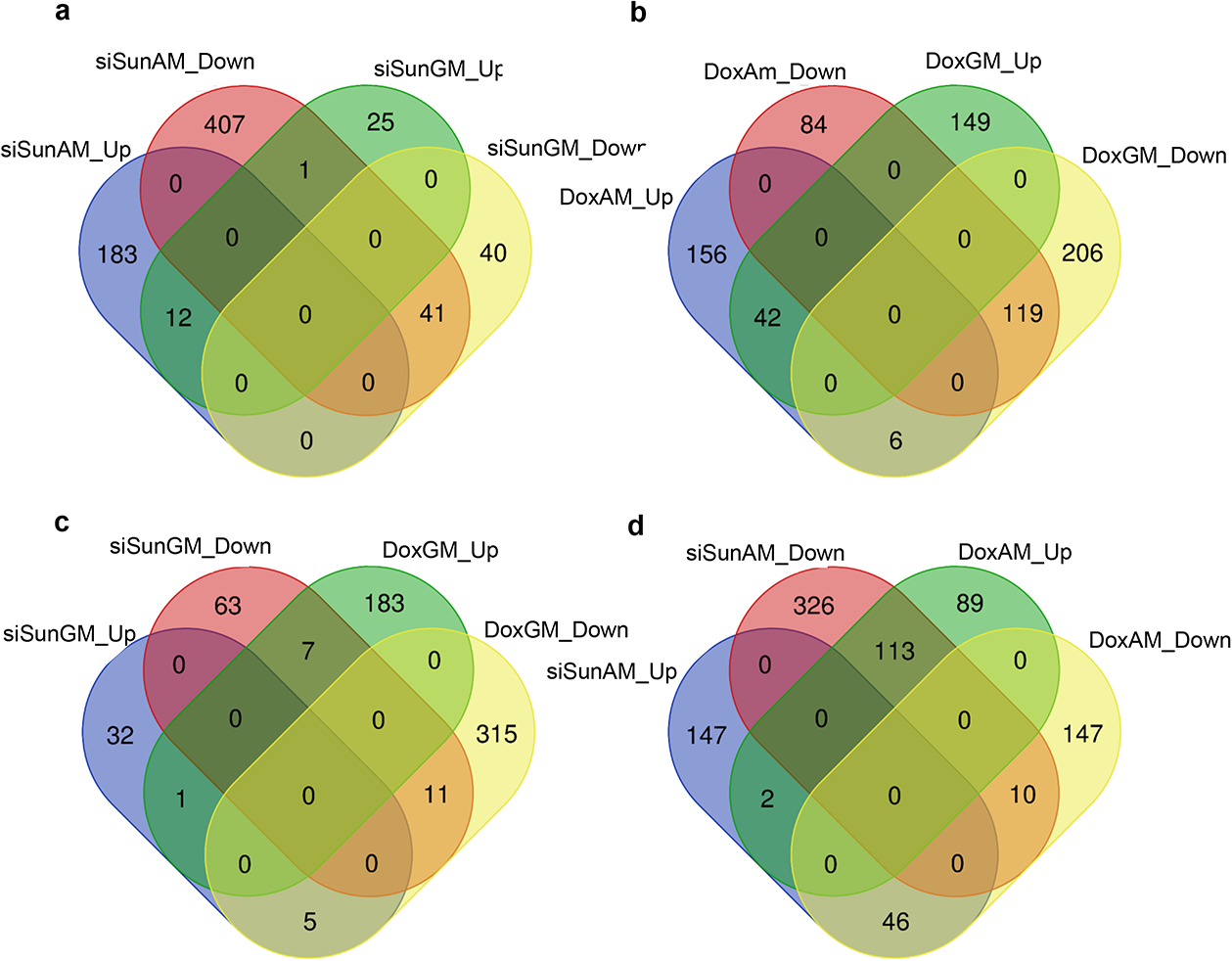
Venn Diagrams of RNA-Seq analysis. **a** Venn diagram depicting gene expression overlap for genes up or down regulated in siSun compared to siCntl during growth in GM or AM. **b** Venn diagram depicting gene expression overlap for genes up or down regulated in GM+Dox or AM+Dox compared to GM or AM respectivly. **c** Comparrison of siSun GM up and down regulated genes compared to siCntl GM and GM+Dox up and down regulated genes compared to GM. **d** Comparrison of siSun AM up and down regulated genes compared to siCntl AM and AM+Dox up and down regulated genes compared to AM.

**Fig. S10.**
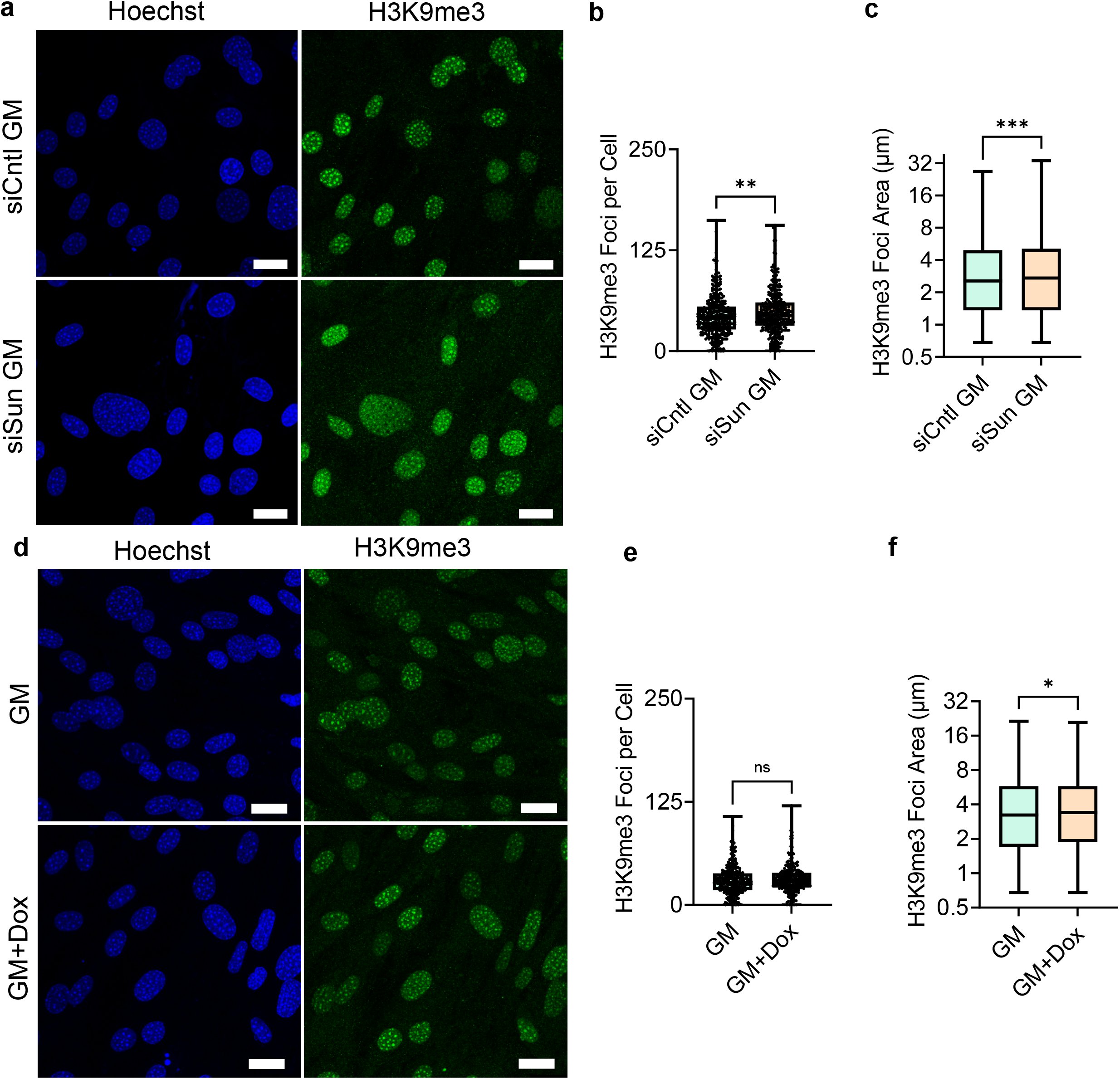
Image analysis of H3K9me3 in GM during Sun1/2 depletion and +Dox treatment. **a** Representative images of siCntl and siSun treated cells grown in growth media staining for H3K9me3 (green) and Hoescht (blue). **b** H3K9me3 foci count per cell in siSun cells compared to siCntl cells in growth media increased by 9% (n = 338, P < 0.01). **c** H3K9me3 foci area increased by 7% in siSun cells compared to siCntl in growth media (n = 14560, P < 0.001). **d** Representative confocal imaging of H3K9me3 (green) and DNA (Blue) in growth media. **e** Analysis of H3K9me3 foci count per cell in the doxycycline treatment group in growth media showed no significant changes in foci count per cell (n= 246). **f** H3K9me3 foci area increased by 5% in doxycycline treatment group in growth media (n = 7350, P < 0.05). Group analysis was made using Two-Tailed Students T-Test. **** P < 0.0001.

**Fig. S11.**
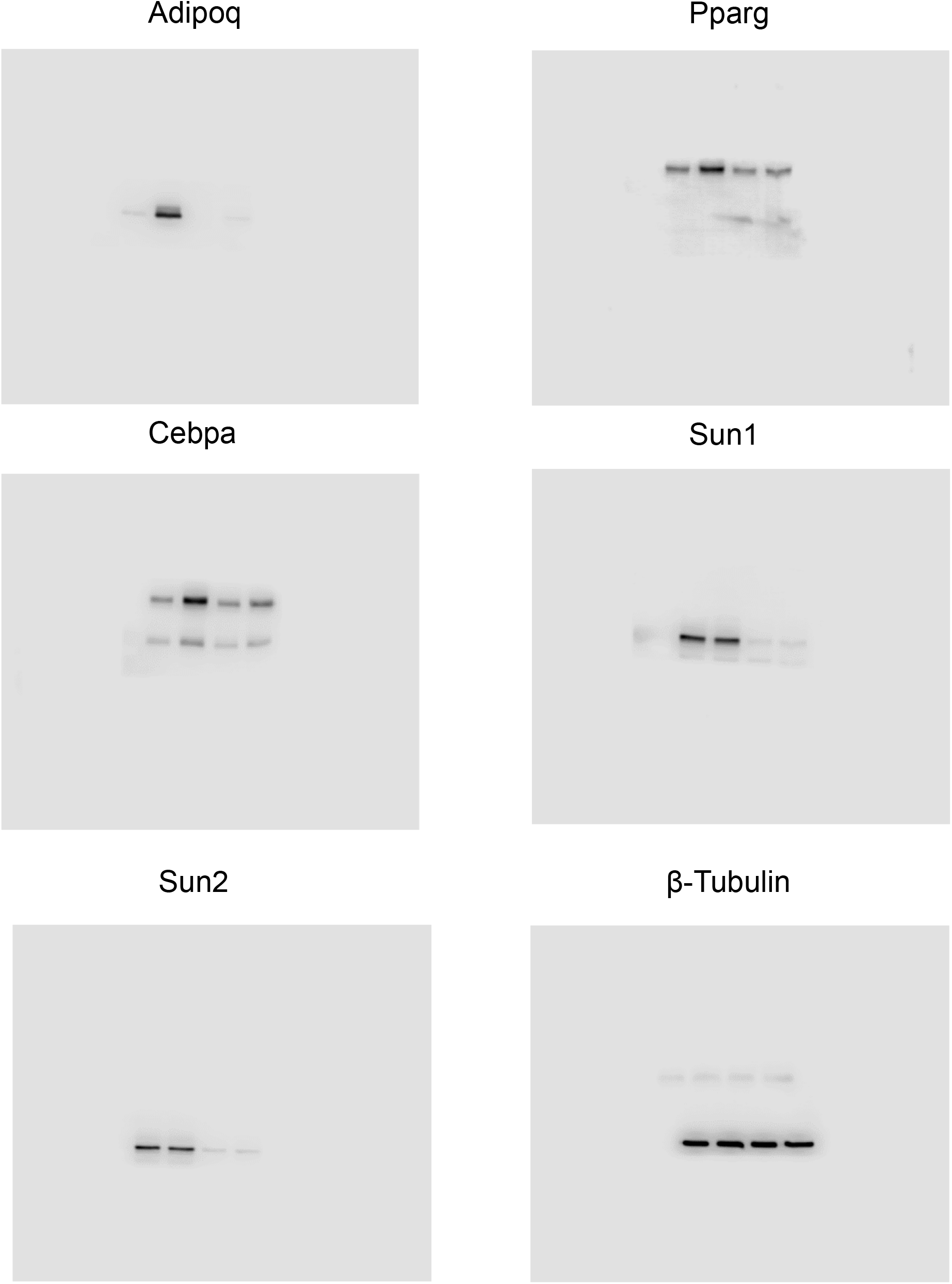
Unprocessed blots for figure 3.

**Fig. S12.**
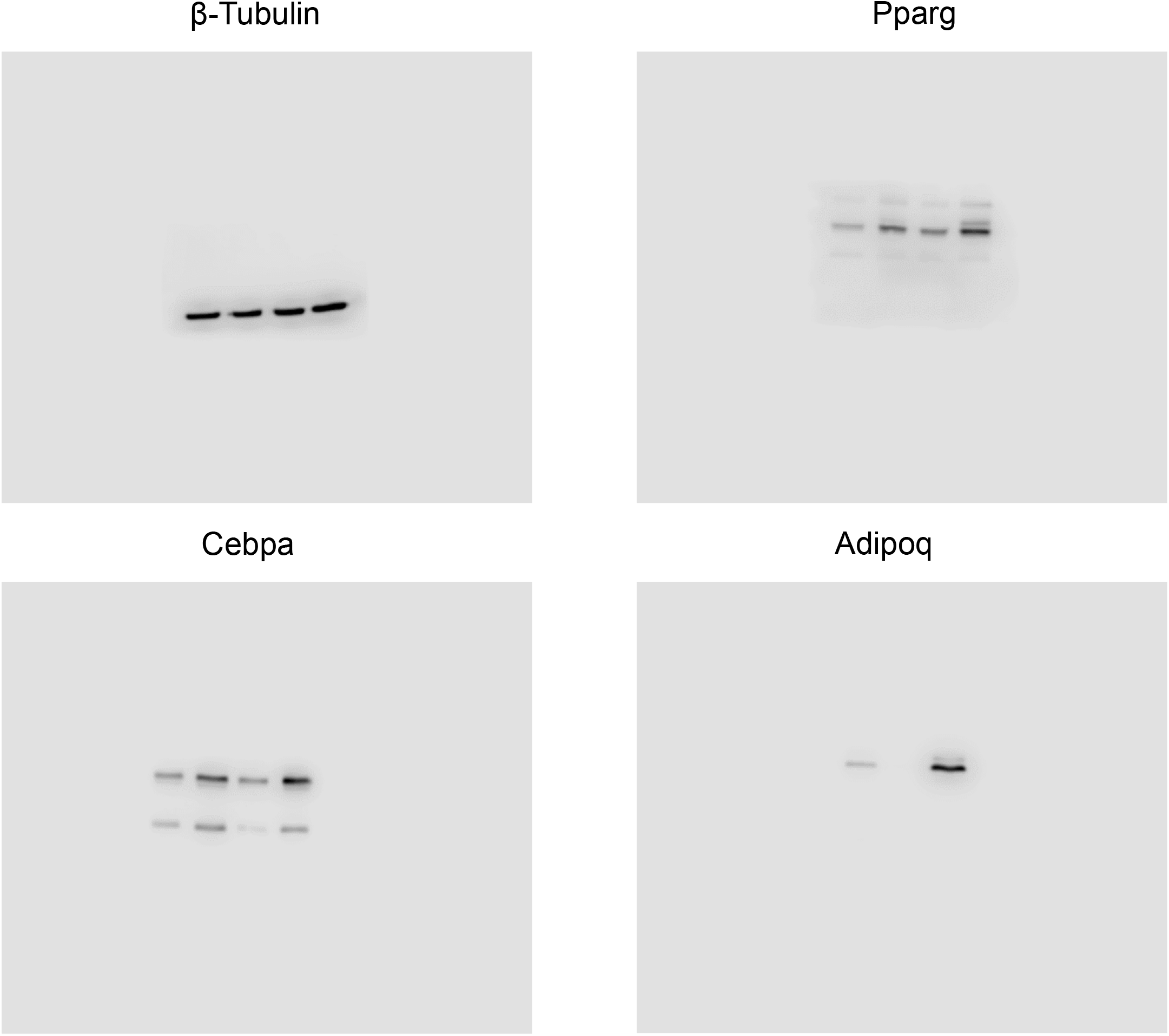
Unprocessed blots for figure 5.

**Fig. S13.**
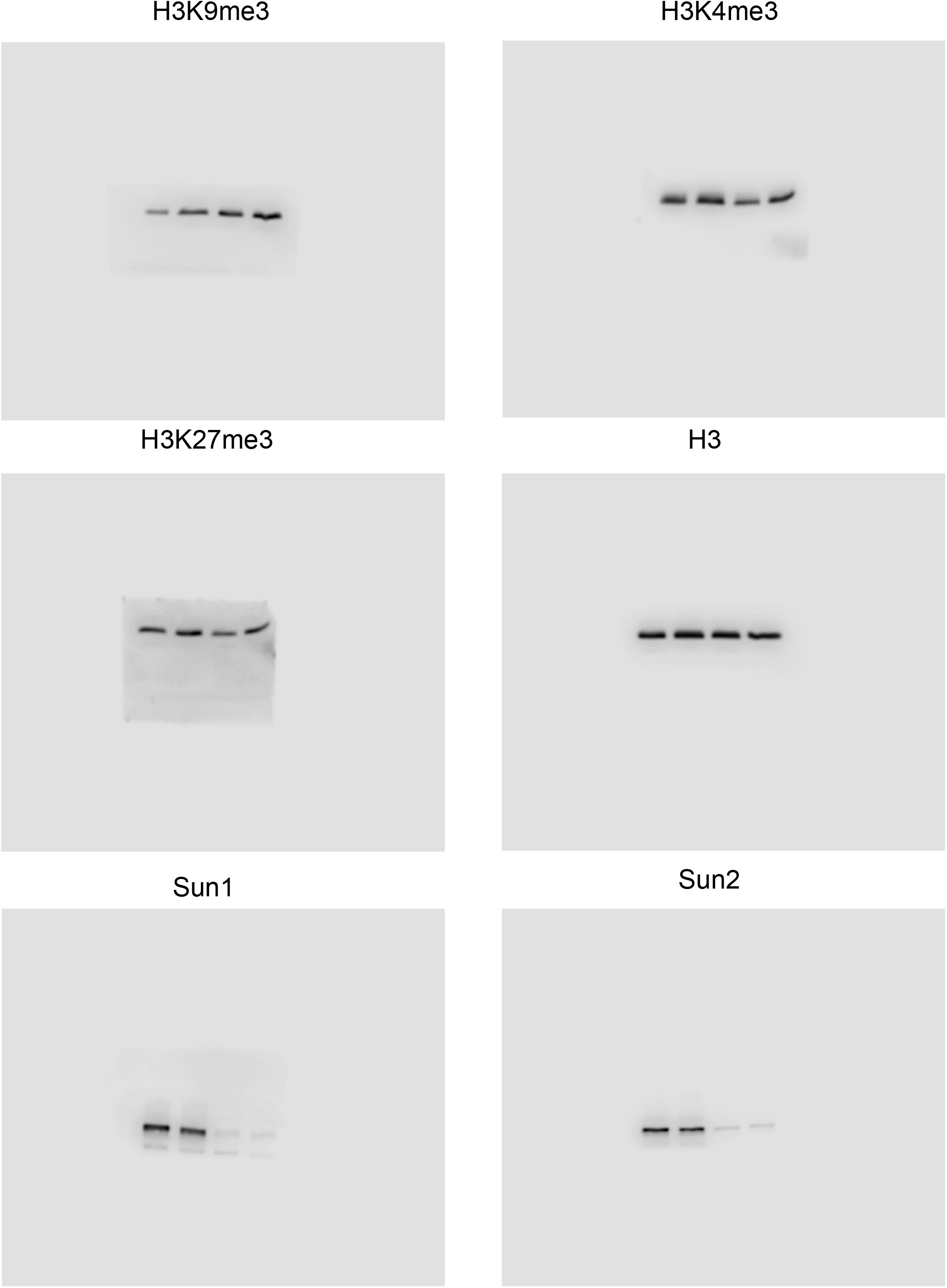
Unprocessed blots for figure 7.

**Fig. S14.**
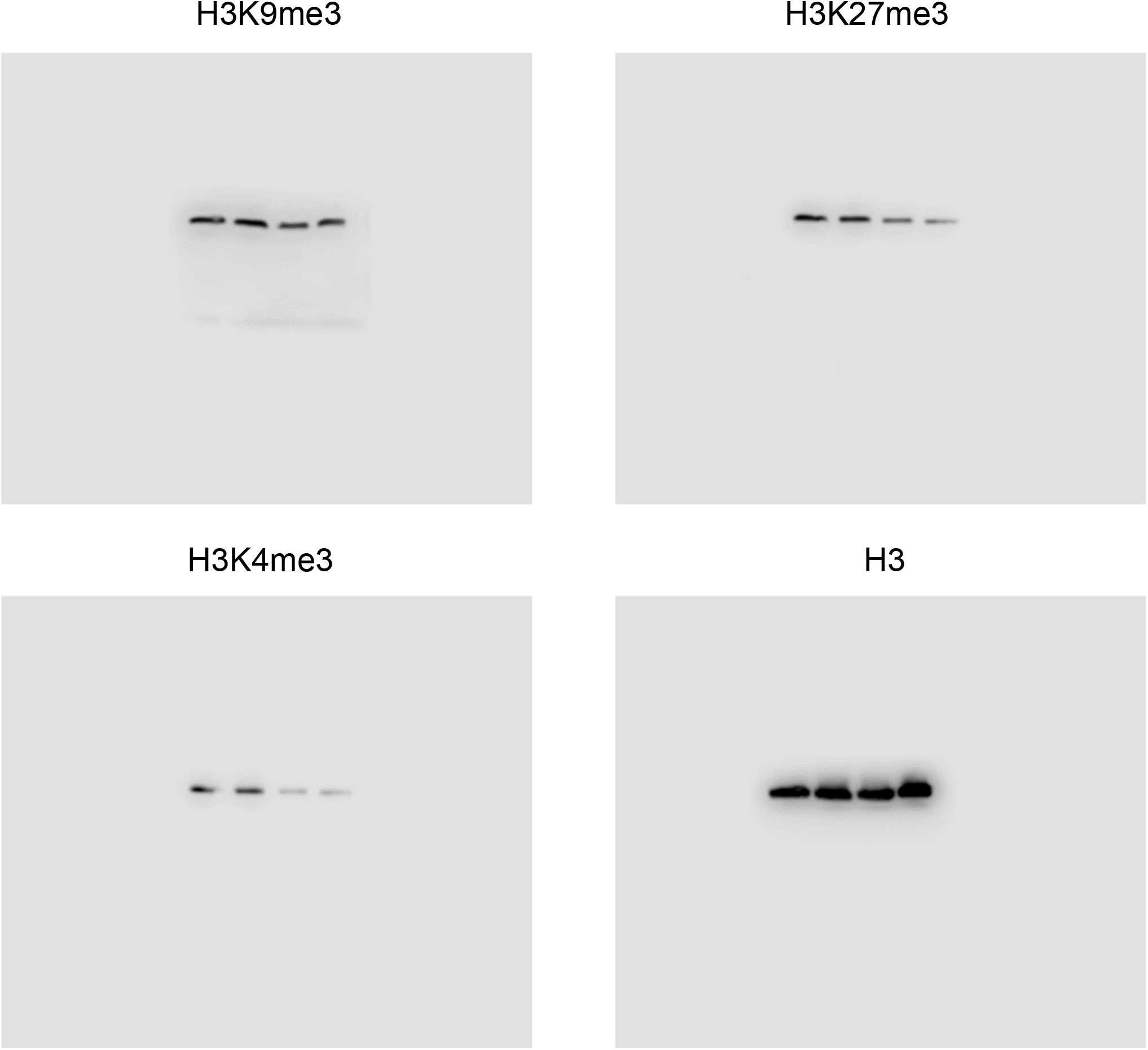
Unprocessed blots for figure 8.

**Table S1.**
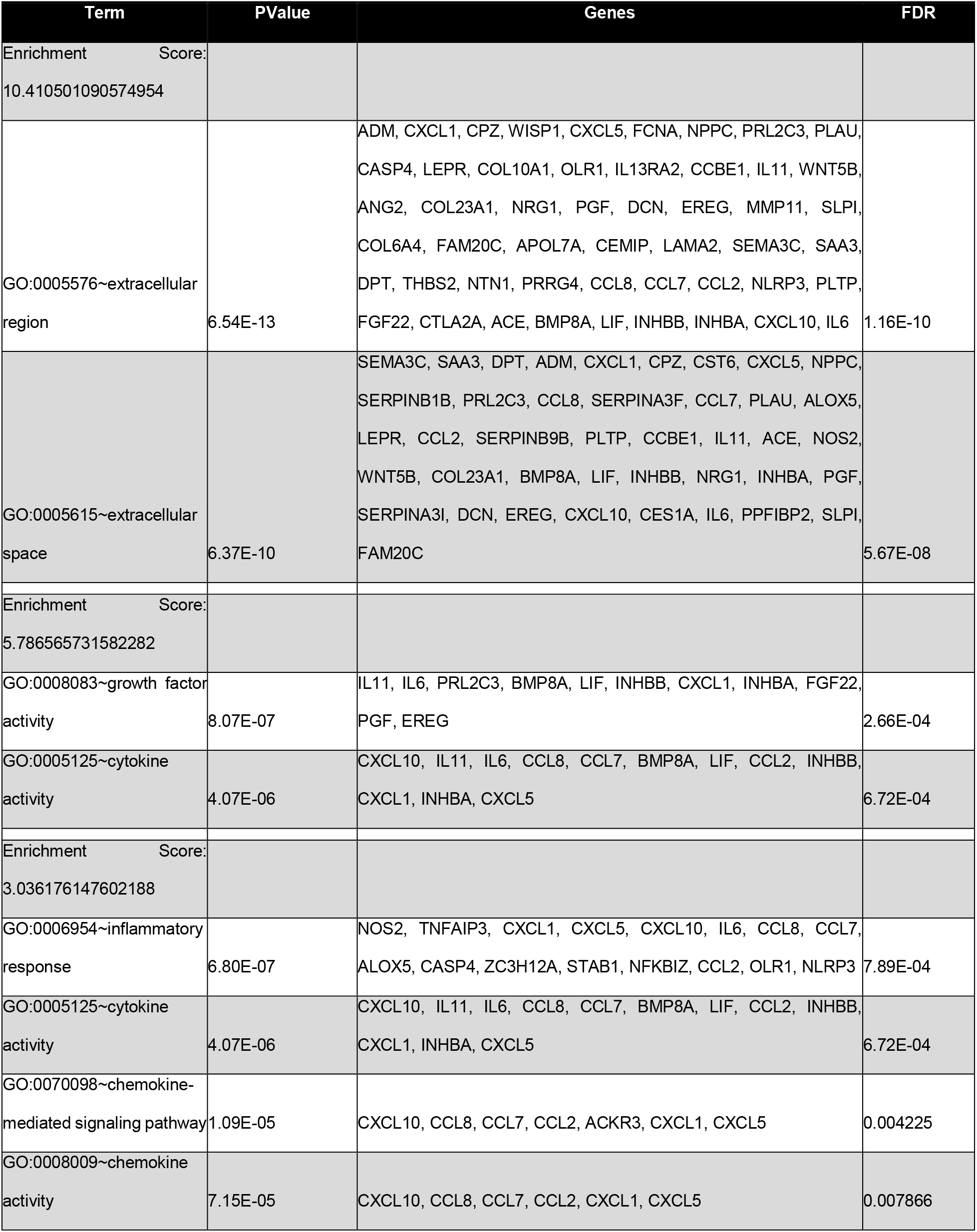
DAVID analysis of genes up regulated in siSun AM vs siCntl AM comparison.

**Table S2.**
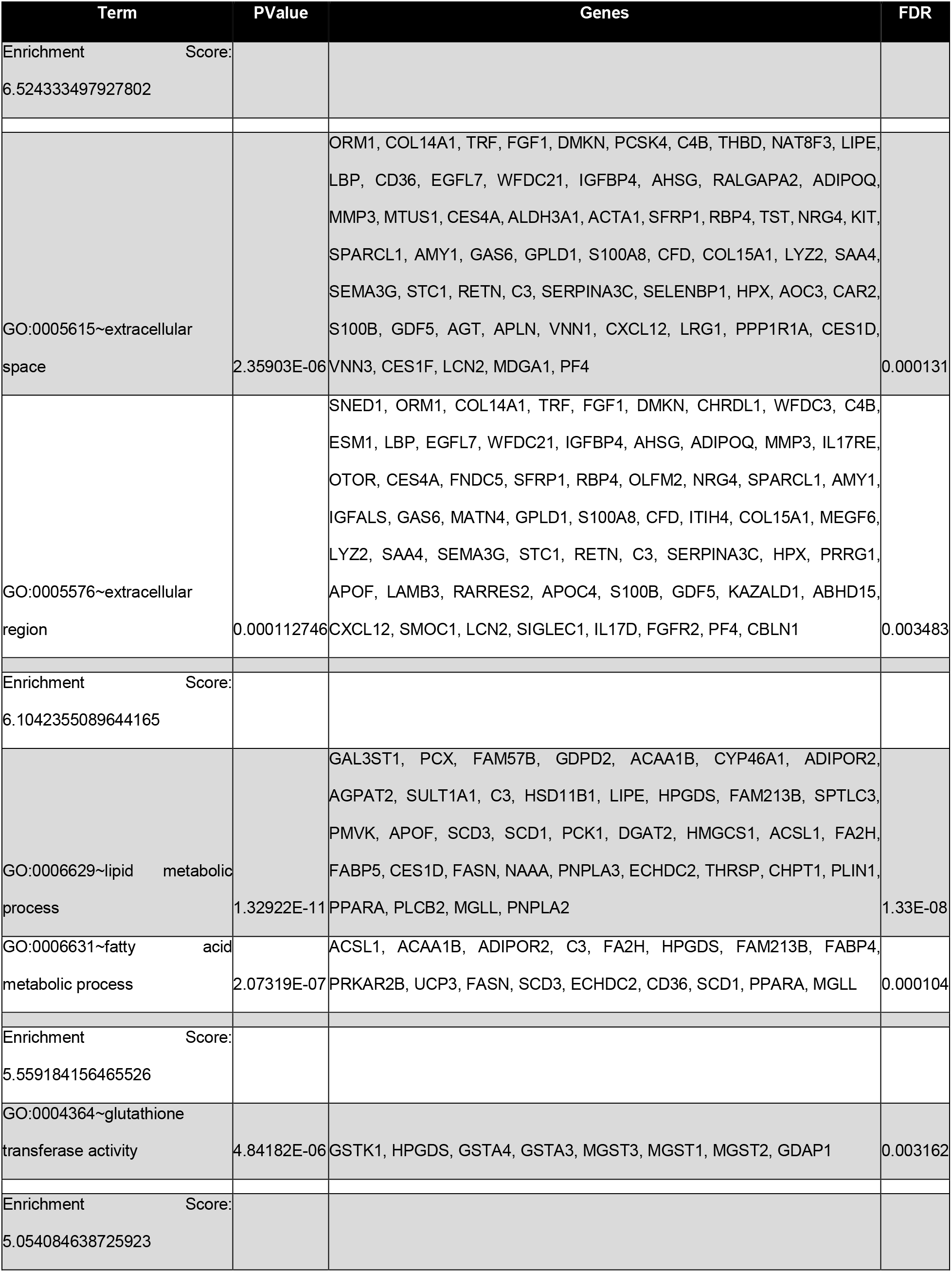
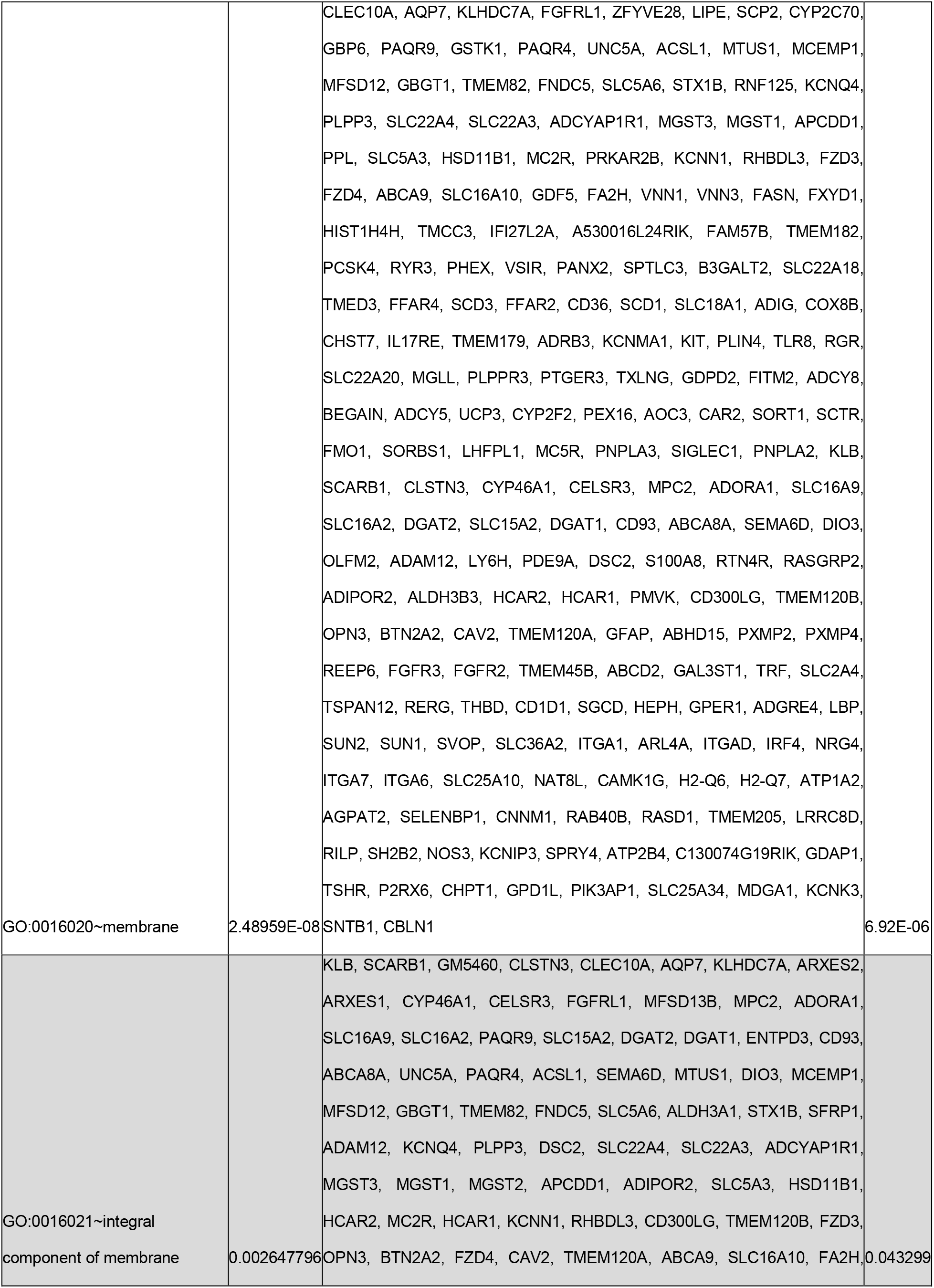
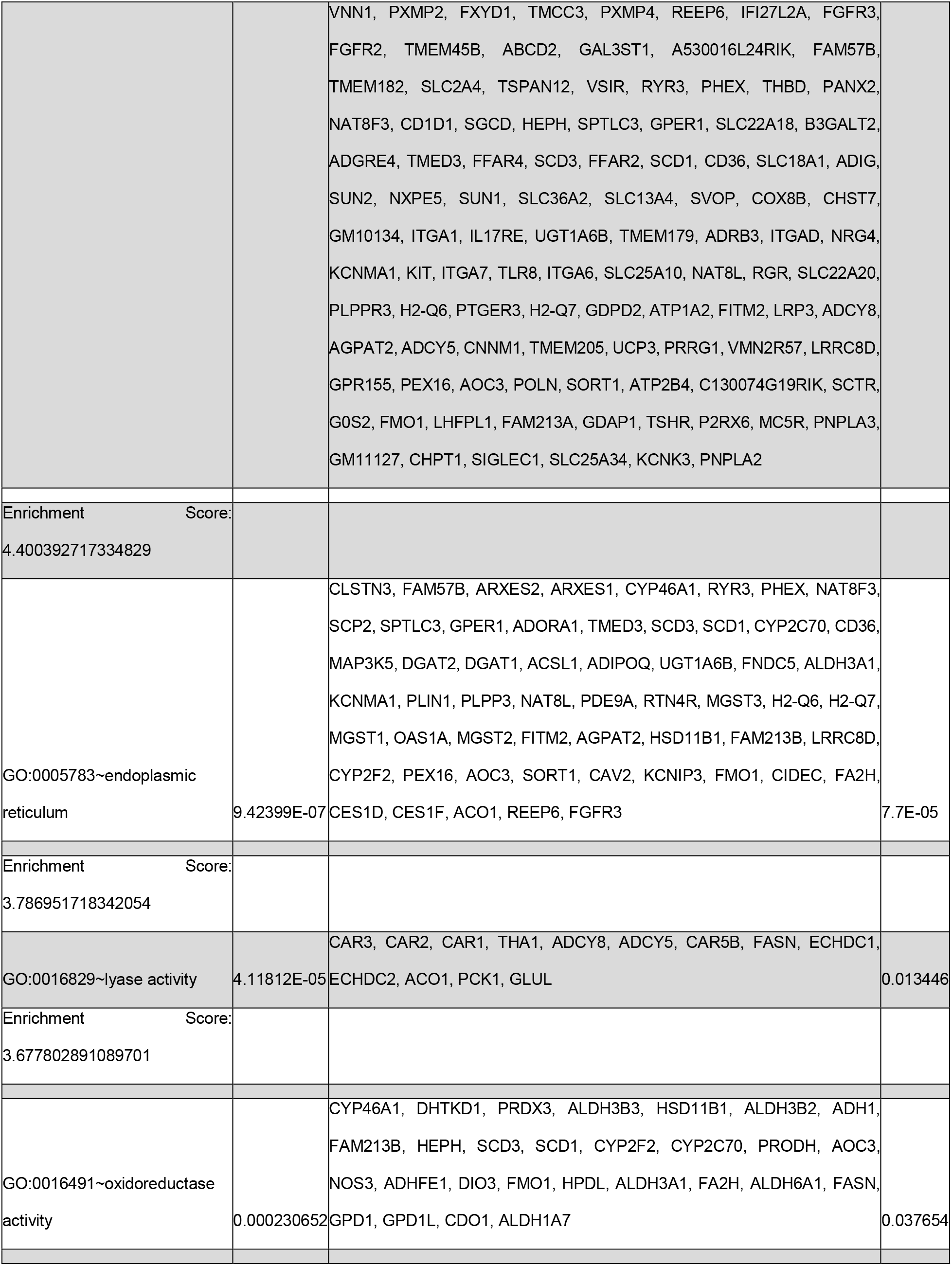
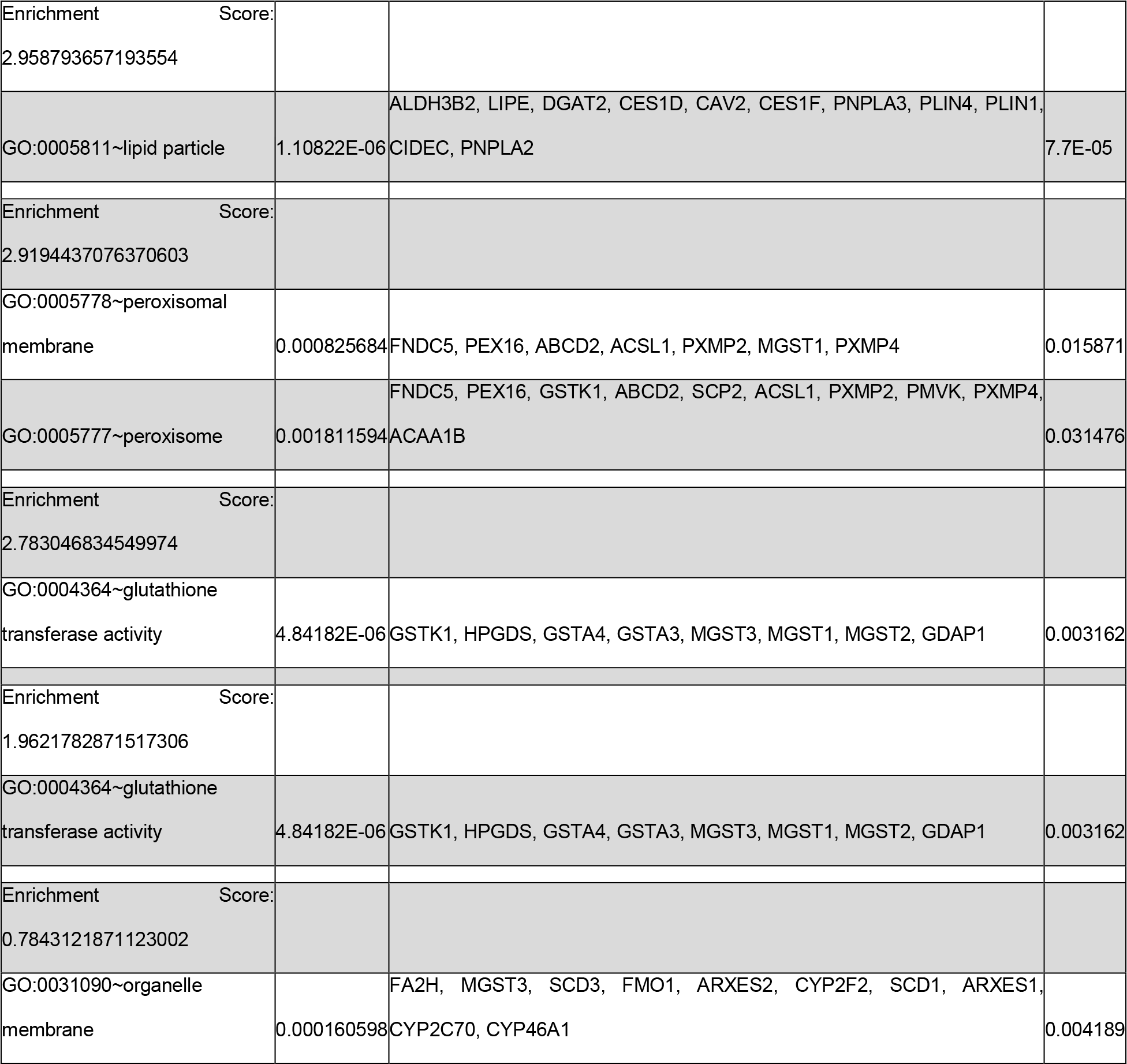
DAVID analysis of genes down regulated in siSun AM vs siCntl AM comparison.

**Table S3.**
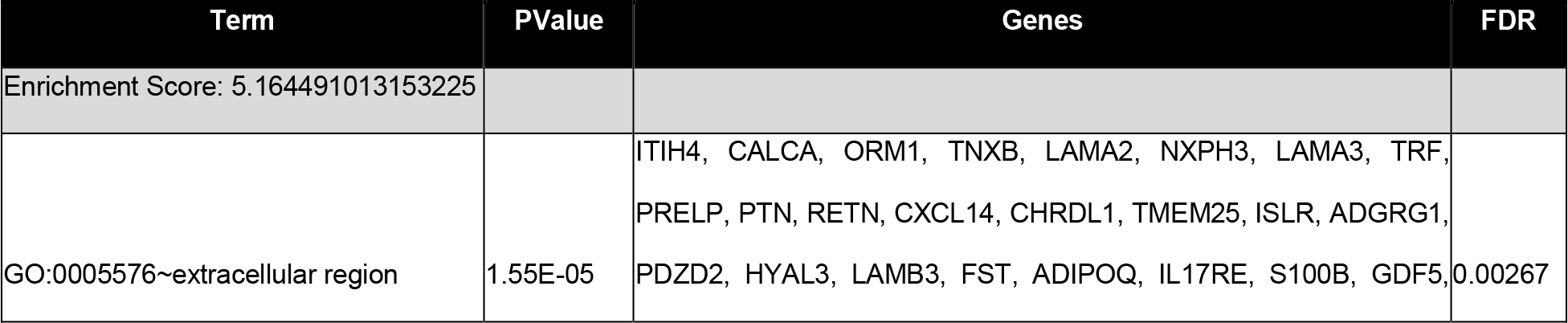
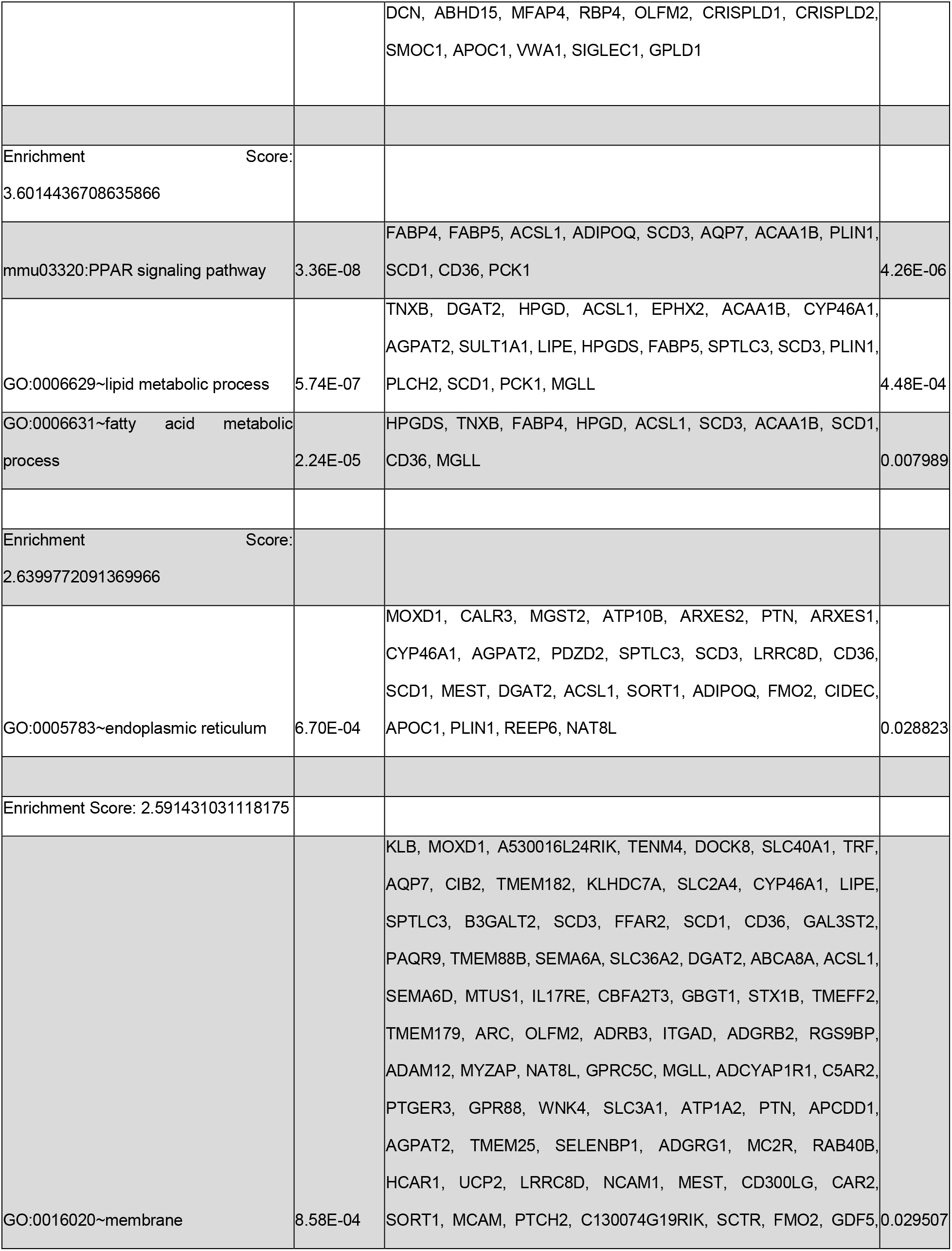
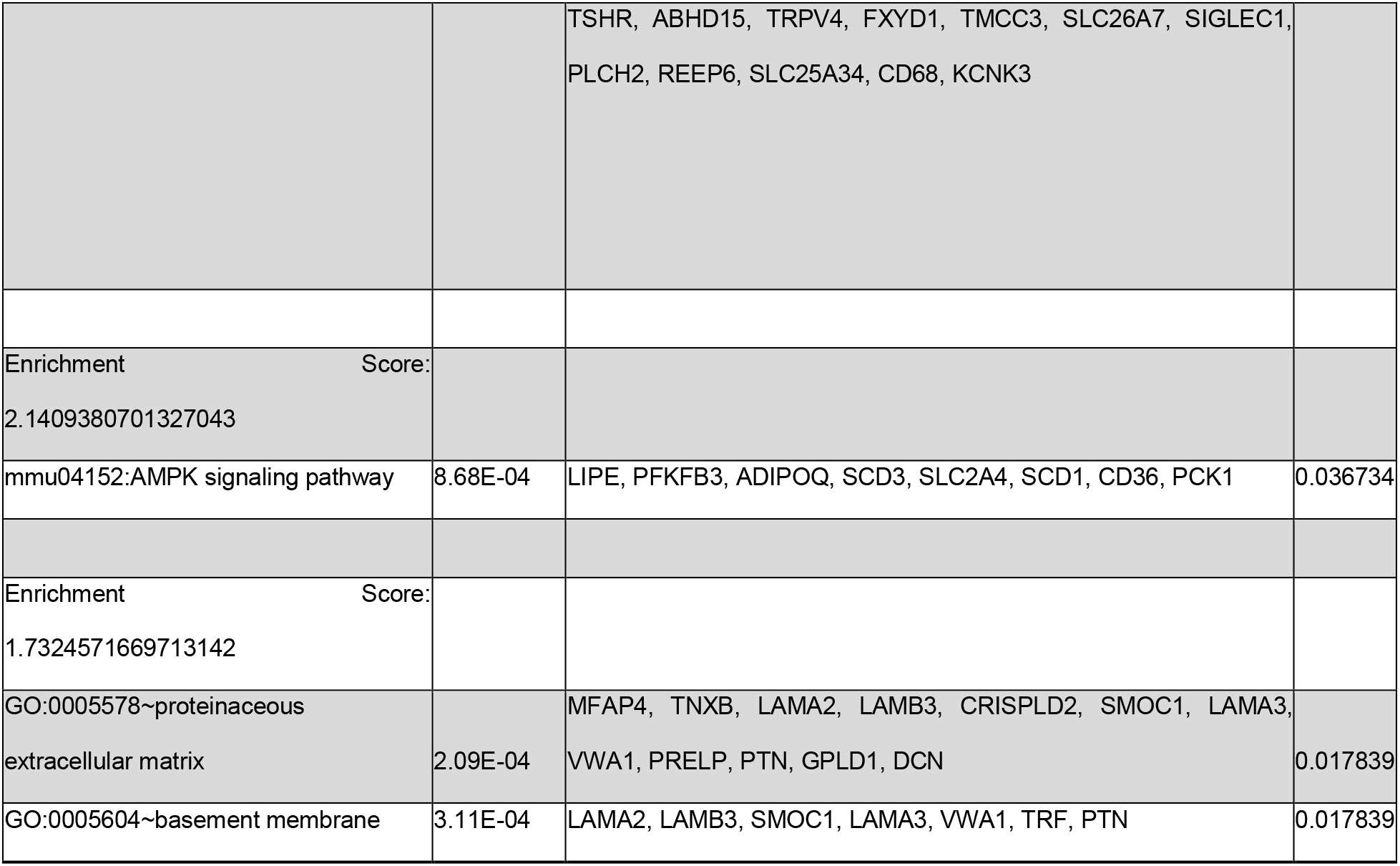
DAVID analysis of genes up regulated in AM vs AM+Dox comparison.

**Table S4.** DAVID analysis of genes down regulated in AM vs AM+Dox comparison.

| Term | PValue | Genes | FDR |
| --- | --- | --- | --- |
| Enrichment Score:<br>16.442538003224215 |  |  |  |
| GO:0005615--extracellular space | 1.6417017461634274E-21 | TINAGL1, SERPINE1, HP, LAMC2, CXCL1, VLDLR, CLU, CXCL5, WISP2, SPN, C4B, LGALS3, FGF7, TIMP3, CAR6, LMCD1, CAMP, CCBE1, EDN1, VWF, GPX3, MMP3, MMP8, EREG, BCAN, MMP13, SLPI, OAS3, FAM20C, ACPP, SAA3, SAA4, TNFRSF11B, LTBP2, LRP8, KNG1, RNASET2A, C3, SERPINB1B, SERPINA3F, CCL7, PTPRZ1, STC2, CCL2, SERPINB9B, NOS2, TNFRSF9, LIF, SERPINA3M, SERPINA3N, SERPINA3G, CP, SERPINA3I, CXCL10, IL6, GDNF, CES1G, CLEC2G, LCN2, PTX3 | 3.2013184050186834E-19 |
| GO:0005576--extracellular region | 5.692614007016396E-17 | TINAGL1, SERPINE1, HP, F13A1, LAMC2, CXCL1, ISM1, CLU, CXCL5, WISP2, C4B, LGALS3, FGF7, CASP4, TIMP3, CAR6, CAMP, CCBE1, EDN1, VWF, GPX3, MMP3, MMP8, MMP10, EREG, MMP12, BCAN, MMP13, SLPI, ANGPTL6, FAM20C, VSTM2A, ACPP, SAA3, SAA4, TNFRSF11B, LTBP2, NTN1, LRP8, KNG1, 5430419D17RIK, C3, CCL7, PTPRZ1, STC2, CCL2, CDSN, LIF, SERPINA3M, SERPINA3N, | 5.5502986568409865E-15 |
|  |  | CP, CXCL10, IL6, GDNF, LCN2,<br>PTX3, NOTUM, HPSE |  |
| Enrichment Score:<br>4.585972071952109 |  |  |  |
| Term | PValue | Genes | FDR |
| GO:0034097~response to cytokine | 1.8576718254593303E-5 | SERPINA3F, SERPINA3M,<br>SERPINA3N, PTGS2,<br>SERPINA3G, OSMR,<br>SERPINA3I, RELB | 0.004102358614556021 |
| GO:0010466~negative regulation of peptidase activity | 2.6176895538567044E-5 | SERPINB1B, SERPINA3F,<br>SLPI, SERPINE1, TIMP3,<br>SERPINA3M, SERPINA3N,<br>SERPINA3G, KNG1 | 0.004336764545185795 |
| GO:0004867~serine-type endopeptidase inhibitor activity | 3.239574315699336E-5 | SERPINB1B, SERPINA3F,<br>SLPI, SERPINE1, SERPINB9B,<br>SERPINA3M, SERPINA3N,<br>SERPINA3G, SERPINA3I | 0.006398159273506189 |
| GO:0030414~peptidase inhibitor activity | 3.239574315699336E-5 | SERPINB1B, SERPINA3F,<br>SLPI, SERPINE1, TIMP3,<br>SERPINA3M, SERPINA3N,<br>SERPINA3G, KNG1 | 0.006398159273506189 |
| GO:0043434~response to peptide hormone | 9.897662481969958E-5 | SERPINA3F, STC2,<br>SERPINA3M, SERPINA3N,<br>SERPINA3G, SERPINA3I,<br>EREG | 0.013114402788610195 |
| Enrichment Score:<br>4.127262509244919 |  |  |  |
| Term | PValue | Genes | FDR |
| GO:0071347~cellular response to interleukin-1 | 1.531896347360444E-6 | IL6, EDN1, CCL7, SAA3, ZC3H12A, SERPINE1, LCN2, CCL2, CAMP | 6.765875534175294E-4 |
| GO:0071222~cellular response to lipopolysaccharide | 2.966452906842876E-4 | CXCL10, IL6, PLSCR1, NOS2, PLSCR2, ZC3H12A, SERPINE1, LCN2, CCL2, CAMP | 0.03275458417972343 |
| Enrichment Score:<br>3.2278474521875653 |  |  |  |
| Term | PValue | Genes | FDR |
| GO:0002376~immune system process | 1.5545629792378633E-4 | CD74, H2-Q7, H2-K1, HP, AHR, SERPINA3G, C3, LGALS3, SLPI, OAS3, CASP4, ZC3H12A, LCN2, TLR2 | 0.01872541770445608 |
| Enrichment Score:<br>3.0652006117499124 |  |  |  |
| Term | PValue | Genes | FDR |
| GO:0005578~proteinaceous extracellular matrix | 7.86993046157755E-8 | CCBE1, VWF, MMP3, LTBP2, LAMC2, TNFRSF11B, MMP8, NTN1, MMP10, WISP2, MMP12, BCAN, LGALS3, MMP13, PTPRZ1, TIMP3, HPSE | 5.1154548000254075E-6 |
| GO:0031012~extracellular matrix | 1.0904983267609749E-6 | TINAGL1, VWF, SERPINE1, MMP3, LTBP2, MMP8, CLU, MMP10, MMP12, LGALS3, PLSCR1, MMP13, SLPI, TIMP3, LMCD1 | 5.100221264313128E-5 |
| Enrichment Score:<br>2.605301978949154 |  |  |  |
| Term | PValue | Genes | FDR |
| GO:0006955~immune response | 1.9461300459952305E-8 | CD74, TINAGL1, TNFRSF9, H2-K1, LIF, CXCL1, TNFRSF11B, CXCL5, CXCL10, IL6, PLSCR1, CCL7, SLPI, OAS3, CCL2, SH2D6, TLR2 | 1.2893111554718401E-5 |

